# A Unifying Principle for the Functional Organization of Visual Cortex

**DOI:** 10.1101/2023.05.18.541361

**Authors:** Eshed Margalit, Hyodong Lee, Dawn Finzi, James J. DiCarlo, Kalanit Grill-Spector, Daniel L. K. Yamins

## Abstract

A key feature of many cortical systems is functional organization: the arrangement of neurons with specific functional properties in characteristic spatial patterns across the cortical surface. However, the principles underlying the emergence and utility of functional organization are poorly understood. Here we develop the Topographic Deep Artificial Neural Network (TDANN), the first unified model to accurately predict the functional organization of multiple cortical areas in the primate visual system. We analyze the key factors responsible for the TDANN’s success and find that it strikes a balance between two specific objectives: achieving a task-general sensory representation that is self-supervised, and maximizing the smoothness of responses across the cortical sheet according to a metric that scales relative to cortical surface area. In turn, the representations learned by the TDANN are lower dimensional and more brain-like than those in models that lack a spatial smoothness constraint. Finally, we provide evidence that the TDANN’s functional organization balances performance with inter-area connection length, and use the resulting models for a proof-of-principle optimization of cortical prosthetic design. Our results thus offer a unified principle for understanding functional organization and a novel view of the functional role of the visual system in particular.

## Introduction

Neurons in sensory cortical systems support two kinds of measurements: their response patterns as a function of stimulus input and their spatial arrangement across the cortical surface. The confluence of these observations is referred to as *functional organization*, the reproducible spatial arrangement of neurons within a cortical area according to their response properties. Functional organization is among the most ubiquitous of neuroscience findings, appearing in the topographic maps of the visual system [1], and in auditory [2], parietal [3], sensorimotor [4], and entorhinal areas [5, 6]. These organized structures anchor our understanding of cortical development, function, and dysfunction, yet it remains a mystery what processes govern their emergence, and what computational function they serve.

Any theory of functional organization must explain both neuronal response properties and the physical arrangement of neurons within a cortical area. Furthermore, a *unified* theory should account for the observed functional organization in multiple cortical areas. Prior computational models of the organization within single cortical areas have been developed [7, 8, 9, 10, 11, 12, 13, 14, 15, 16, 17, 18, 19, 20, 21, 22], but these approaches do not generalize to multiple cortical areas. Moreover, many of these models operate from a hand-crafted set of stimulus features, and thus cannot explain how neuronal response properties are learned from realistic sensory inputs. On the other hand, deep artificial neural networks (DANNs) trained with large quantities of naturalistic data are increasingly being used to model neuronal responses in regions responsible for vision, audition, and language processing [23, 24, 25, 26, 27, 28, 29, 30, 31]. However, standard DANNs impose no spatial arrangement among model units that differ in their stimulus tuning, and thus cannot explain the observed organization of neurons across the cortical surface.

Here, we introduce the Topographic Deep Artificial Neural Network (TDANN), a unified framework for predicting functional organization in sensory systems. The TDANN implements the hypothesis that neural systems are optimized to address two key goals: they must support ecologically-relevant behaviors by producing useful neural representations [32], and they must do so in a biophysically efficient manner, using as few resources as possible. A critical component of biophysical efficiency is the minimization of neuronal wiring length, which is theorized to result in the smooth topographic organization observed in many cortical areas [33, 19, 18]. The TDANN begins with a standard DANN and spatially augments it by embedding each layer’s units in a two-dimensional simulated cortical sheet. The TDANN then optimizes a *composite objective function* with two components: a functional objective that drives the learning of useful representations, and a spatial constraint that encourages efficiency with smooth response patterns across the simulated cortical sheet. We test this framework in the primate ventral visual stream, a cortical system in which functional organization has been extensively documented.

The ventral stream is a hierarchical series of cortical areas that support visual recognition, beginning with primary visual cortex (V1) and ascending through intermediate areas (e.g., V4) to high-level regions: inferotemporal (IT) cortex in macaques and ventral temporal cortex (VTC) in humans. Well-known neuronal response properties in V1 include tuning to edge orientation [1, 34, 35], spatial frequency [36], and color [37, 38]. These response properties are coupled with topographic signatures: orientation preferences form a smooth cortical map with pinwheel-like discontinuities [39, 40, 41, 42, 43]; spatial frequency tuning is organized in a quasi-periodic map with isolated low-frequency domains [42, 43, 44]; and color-preferring neurons cluster in punctate blobs [38] across the V1 surface. Higher-level regions such as primate IT [45, 46, 47, 48] and the analogous human VTC contain neurons with stronger responses for items of specific categories vs. others (e.g., faces vs non-faces), a property known as *category selectivity*. A core characteristic of functional organization in IT [48, 49] and VTC [50, 51, 52, 53, 54, 55, 56] is that neurons selective for certain ecologically-relevant categories – including faces, places, limbs, and visual wordforms – cluster into spatial patches, with characteristic patch sizes, counts, and relative inter-patch distances.

We find that the TDANN reproduces the functional organization of the ventral stream, including smooth orientation maps with pinwheels in an earlier model layer, and category-selective patches in a later layer that match the number, size, and relative geometry of patches in human VTC. To understand the principles underlying the emergence of the ventral stream’s functional organization, we then test which specific functional and spatial constraints of the TDANN are critical to the TDANN’s success by insantiating alternative models and measuring their capacity to predict neural data. We find that the specific combination of task and spatial objectives that best matches the functional organization of the ventral stream also makes learned representations more brain-like by constraining their intrinsic dimensionality. The TDANN learns these representations while minimizing the network’s inter-layer wiring length, suggesting that brain-like functional organization effectively balances performance with metabolic costs.

Finally, because the the TDANN accurately predicts the functional organization of the ventral stream, it provides an exciting new platform for simulating experiments that are challenging to implement empirically. As a proof of principle, we perform *in silico* experiments simulating the effect of cortical microstimulation devices that vary in their spatial precision and cortical coverage. Taken together, our results show that the TDANN serves both as a unified explanation for the functional organization of the visual system and as a platform to fuel discovery in neuroscience.

## Results

### Instantiating models that balance task performance with spatial smoothness

Building on optimization-based approaches in computational neuroscience [57, 58], we seek a model architecture and objective function that generate a neural network which matches the neuronal responses and topography of the primate ventral visual stream.

Because standard DANNs have no within-area spatial structure beyond retinotopy, we must augment their architecture to model spatial topography. Specifically, we take the ResNet-18 architecture [59], a DANN that achieves strong object recognition performance and accurate prediction of neuronal responses throughout the ventral visual stream [30], and augment it by embedding the units of each convolutional layer into a two-dimensional simulated cortical sheet (Figure 1a). Given that neurons in visual cortex are organized retinotopically at birth [60], we assign model unit positions retinotopically, such that units responding to similar regions of the input images are nearby in the simulated cortical sheet. Then, prior to training, unit positions are locally shuffled to circumvent limitations of weight-shared convolution (see Methods). The size of the simulated cortical sheet in each layer is anchored by estimates of cortical surface area in the human ventral visual stream (Figure 1a). We refer to the resulting model as the *Topographic DANN (TDANN)*.

**Figure 1.**
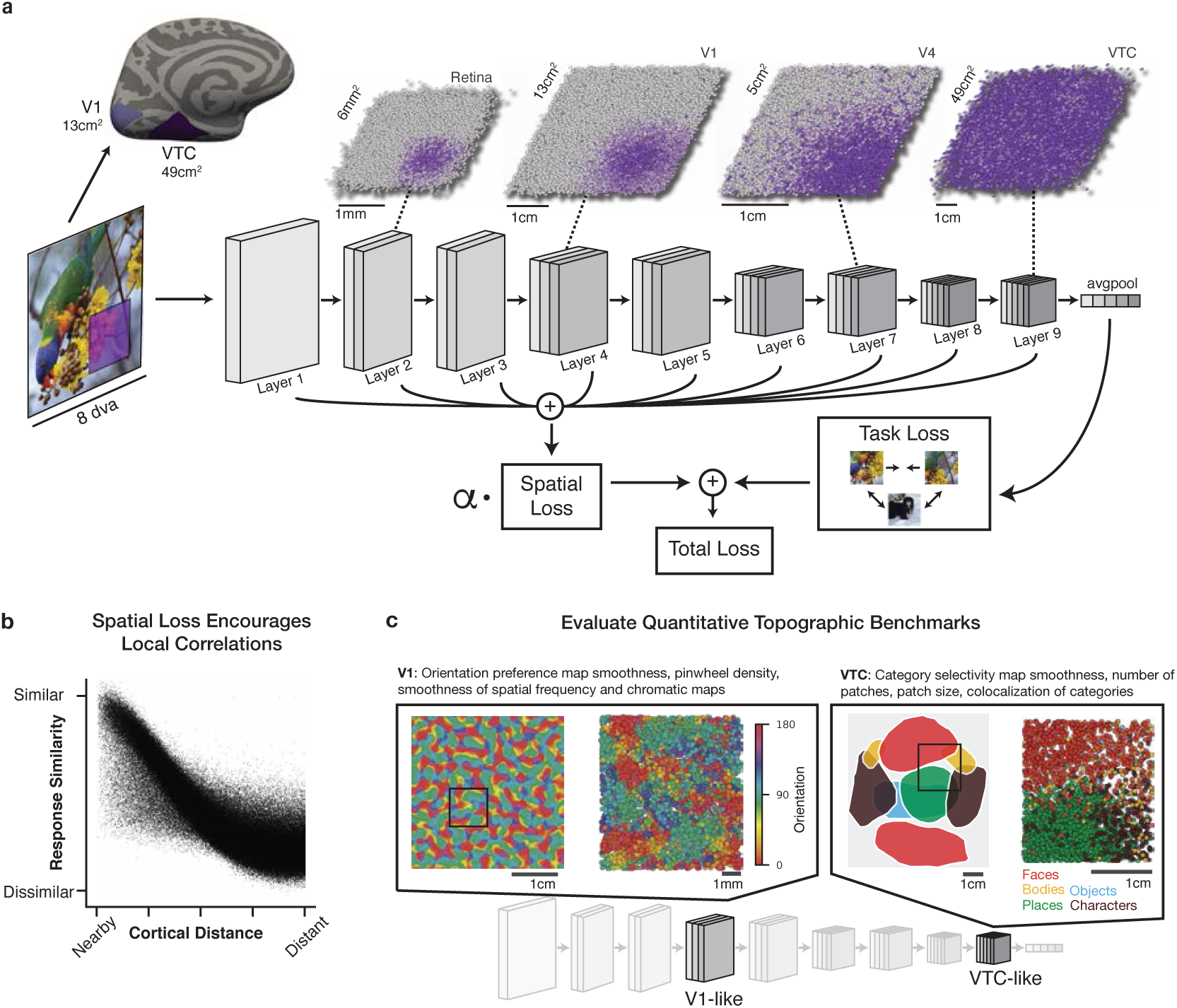
Constructing a unified model of the functional and spatial constraints of ventral visual cortex. **(a)** TDANNs are a family of deep artificial neural networks whose units are assigned positions in a two-dimensional simulated cortical sheet in each layer. Position assignments are retinotopic, such that location in the cortical sheet corresponds to position in the visual field. Each individual dot is a single model unit. The degree of overlap between a unit’s spatial receptive field (RF) and the purple square marked on the input image is indicated by the shade of purple; RFs from gray units do not overlap the marked region at all. The TDANN is trained to minimize the sum of a task loss and a spatial loss (SL). *α* is a free parameter controlling the relative weight of the SL. **(b)** The SL encourages nearby units to develop strong response correlations. Plotted: pairwise similarity of unit responses as a function of pairwise cortical distance in the final layer of a TDANN model; each dot represents one pair of units. **(c)** The TDANN is evaluated on a battery of quantitative benchmarks that measure its correspondence to topographic features throughout the ventral visual stream. Left: orientation preference map in the V1-like TDANN layer (see Figure 2 for details). Right: category selectivity map in the VTC-like layer (see Figure 3 for details).

Having selected the architecture, our goal is to discover the objective whose optimization yields an accurate model of both response properties and their topographic arrangement. The core of the TDANN approach is a composite objective that is a weighted sum of two components: a task objective encouraging the learning of behaviorally-useful functional representations, and a spatial objective driving the emergence of topographic properties. Following recent progress in training neural networks without explicit category labels [61, 62], we use an unsupervised algorithm that performs *contrastive self-supervision*, SimCLR [63], as the task objective. For the spatial loss (SL), we introduce an objective that encourages nearby pairs of units to have more correlated responses than distant pairs of units (Figure 1b, see Methods). The SL is computed separately in each convolutional layer, then summed across layers for each batch of training data:

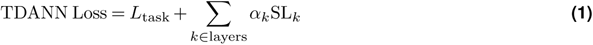

where *α_k_* is the weight of the spatial loss in the *k*th layer, set to *α_k_* = 0.25 for all layers. The TDANN architecture is trained to optimize this objective using conventional back-propagation with stochastic gradient descent.

Training the TDANN on ImageNet [64] resulted in successful minimization of both task and spatial losses (Supplementary Figure S1). We tested if adding the spatial loss interferes with visual representation learning by measuring the model’s object categorization performance with a linear readout. Categorization accuracy was slightly but significantly lower for the TDANN (median across random initialization seeds = 43.9%) than “Task Only” models with no spatial loss (*α* = 0, median = 48.5%; Mann-Whitney *U* = 25, *p* = .008). Despite the modest decrease in categorization performance, adding the spatial loss term had the intended effect: in each layer, the correlation between units’ responses increased with spatial proximity (Supplementary Figure S1c,d). To determine if this learned correlation structure corresponds to brain-like topographic maps, we constructed a battery of quantitative benchmarks comparing model predictions with neural data in primary visual cortex (V1) and ventral temporal cortex (VTC), (Figure 1c). To compare against these benchmarks, we needed to identify the TDANN layers that would be our models of V1 and VTC. As in prior work [28, 25], we find that earlier model layers best predict V1 responses and later layers best predict responses in higher visual cortex (Supplementary Figure S2). Accordingly, we designate the fourth and ninth convolutional layers as the “V1-like” and “VTC-like” layers, respectively.

### The TDANN predicts the functional organization of primary visual cortex

Neurons in primate V1 are organized into maps of preferred stimulus orientation, spatial frequency, and color [38, 43, 65]. Because high-resolution data at the scale necessary to visualize these maps is not available for human V1, we compare the TDANN to macaque V1 data using scale-invariant metrics. We tested if the V1-like TDANN layer captures the functional organization of macaque V1 with three kinds of quantitative benchmarks. First, we evaluate functional correspondence by asking if model units in the TDANN V1-like layer have similar preferred orientations and orientation tuning strengths as neurons in macaque V1. Second, we assay the structure of cortical maps by measuring pairwise similarity of tuning for orientations, spatial frequencies, and colors as a function of cortical distance. Third, we measure the density of pinwheel-like discontinuities in the orientation preference map, a hallmark of V1 functional organization in many species [41, 66]. In addition to the TDANN, we also evaluate four control models on these benchmarks: the *Unoptimized* TDANN, in which model weights and unit positions are left randomly initialized, the *Task Only* variant in which *α* = 0, and two kinds of self-organizing maps (SOMs), which have been proposed as models of V1 functional organization [11, 10]. We refer to the traditional SOM in which feature dimensions are manually predetermined (as in Swindale and Bauer [11]), as the Hand-Crafted SOM, and a novel SOM that organizes the output of an AlexNet V1-like layer (inspired by Doshi and Konkle [13], Zhang et al. [12]) as the DNN-SOM.

#### The TDANN matches orientation tuning in V1

We measured orientation tuning strength by presenting a set of oriented sine grating images to the model (Figure 2a), computing a tuning curve for each unit, and calculating the circular variance (CV; lower values for sharper tuning) of each tuning curve. Setting a selectivity threshold of CV < 0.6, we find that the TDANN V1-like layer has a significantly greater proportion of selective units (range across model seeds: [20%, 31%]) than Unoptimized models ([1%, 3%]; Mann-Whitney *U* = 25; *p* = .008, Figure 2b), but fewer than Task Only models ([35%, 50%]; *U* = 25; *p* = .008) or macaque V1 (45%; Supplementary Figure S3c). In contrast, neither the Hand-Crafted SOM nor the DNN-SOM exhibited any units with sharp orientation tuning. We also find that TDANN and Task Only models (but not SOMs or Unoptimized models) show an over-representation of cardinal orientations (0 and 90 degrees) as in macaque V1 [35] (Supplementary Figure S3b, see also Henderson and Serences [67]).

**Figure 2.**
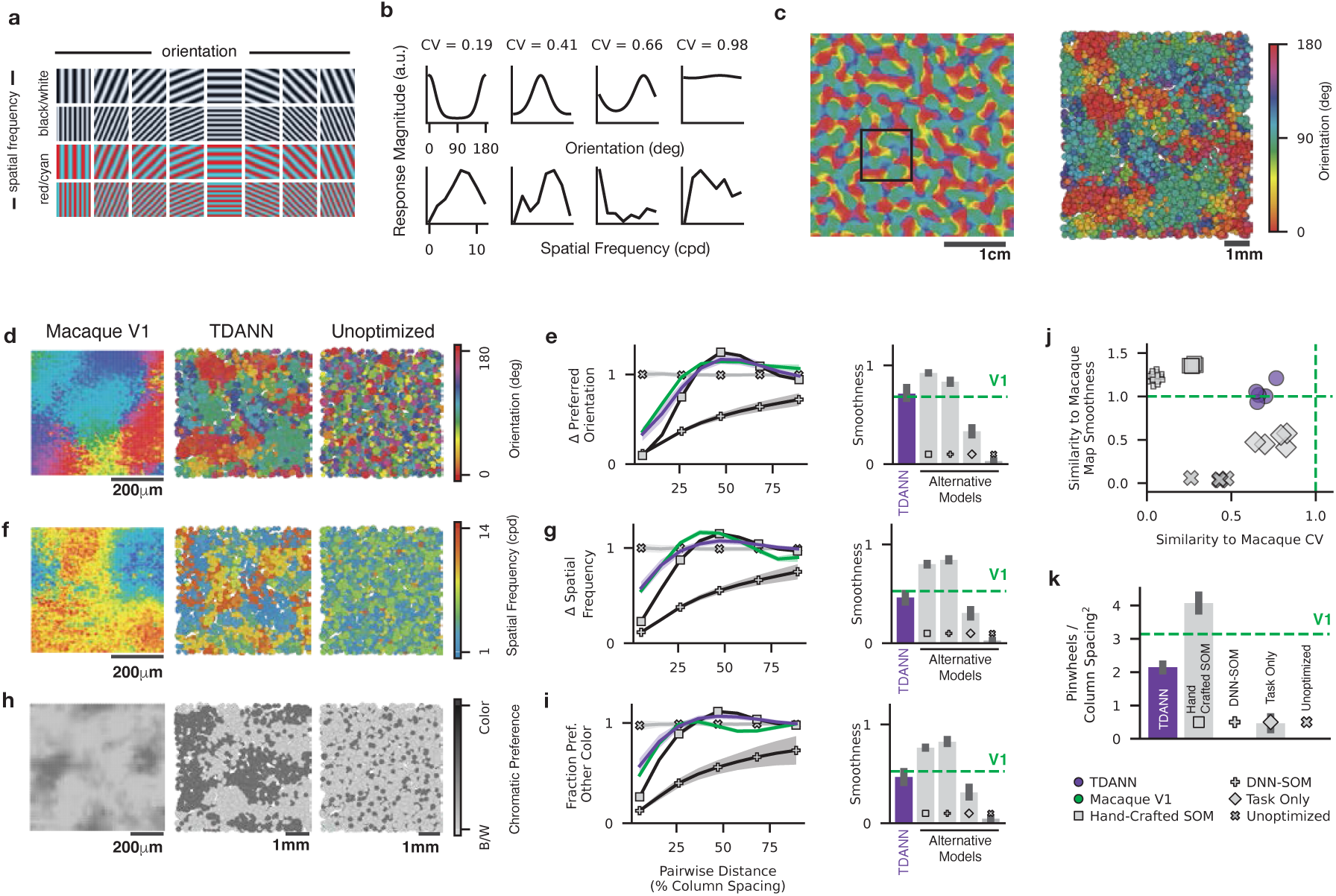
The TDANN reproduces V1-like topography. **(a)** Example sine grating stimuli used to assess tuning for orientation, spatial frequency, and color. **(b)** Orientation tuning curves (top) and spatial frequency tuning curves (bottom) for four example units in the V1-like layer. **(c)** Smoothed orientation preference map (OPM) in the V1-like layer of the TDANN. Box corresponds to inset at right, where individual model units are labeled by their preferred orientation. Results for additional model seeds shown in Supplementary Figure S10. **(d)** OPMs for Macaque V1 (data from Nauhaus et al. [43]), TDANN, and an Unoptimized control model. **(e)** Left: Pairwise difference in preferred orientations as a function of pairwise cortical distance, normalized to the chance level expected by random sampling of pairs. Right: Map smoothness for OPMs in macaque V1 (dashed green line, data from Nauhaus et al. [43]) and four candidate models: the TDANN (purple), the Hand-Crafted self-organizing map (SOM, squares), deep neural network SOM (DNN-SOM, plus signs), and Task Only (diamonds) trained without the spatial term of the loss function. Error bar: 95% CI across random model seeds and sampling of cortical neighborhoods. **(f)** Spatial frequency preference, shown for the same region of the TDANN V1-like layer and macaque V1 as in panel (d). **(g)** Change in preferred spatial frequency as a function of cortical distance, normalized to chance, for macaque V1 and each model type. **(h)** Preference for chromatic stimuli for the same region of the TDANN V1-like layer. Dark-colored dots: stronger responses to chromatic than achromatic gratings. Macaque data: reconstruction of cytochrome oxidase staining data from Livingstone and Hubel [38]. **(i)** Fraction of units differing in their chromatic preference as a function of cortical distance, normalized to chance. **(j)** Similarity of models to the distribution of orientation tuning strengths in macaque V1 (data from Ringach et al. [34]) on the x-axis, and similarity to the smoothness of macaque OPMs (data from Nauhaus et al. [43]) on the y-axis. Multiple markers of the same type indicate different random initial seeds for each model. A value of 1.0 (dashed green) indicates perfect correspondence. **(k)** Density of pinwheels detected in TDANNs, Hand-Crafted SOMs, Task Only models, and Unoptimized models. Error bars: CI across random model seeds. Green: putative macaque V1 pinwheel density.

#### The TDANN predicts the arrangement of orientation-selective V1 neurons

To evaluate whether the TDANN V1-like layer captures the topographic properties of macaque V1, we consider the spatial distribution of orientation-selective units – the orientation preference map (OPM) – and find a smooth progression of preferred orientations that resembles macaque V1 (Figure 2c, d). Following prior work [68, 69, 70], we quantify this structure by measuring the absolute pairwise difference in preferred orientation as a function of cortical distance. In both the TDANN and macaque V1 (data from Nauhaus et al. [43]), we find that nearby units have smaller differences in orientation preference than distant pairs (Figure 2e). In contrast, orientation preference similarity does not vary with cortical distance in Task Only or Unoptimized models, and both the Hand-Crafted and DNN-SOMs exhibit OPMs with abnormally high orientation tuning similarity (Figure 2e, Supplementary Figure S3). We summarize these profiles by computing a *smoothness score* that measures the increase in tuning similarity for nearby unit pairs compared to distant unit pairs. Smoothness of TDANN OPMs ([min, max] across random initialization: [.64, .83]) was consistent with macaque V1 (.68); however, OPMs in the Hand-Crafted SOM ([.92, .92]) and DNN-SOMs ([.81, .86]) were smoother than in macaque V1. In turn, macaque V1 OPMs were smoother than Unoptimized ([.03, .04]) and Task Only ([.28, .39]) models. Jointly comparing each model to macaque V1 orientation tuning strength and OPM smoothness highlights that the TDANN is the only model class that satisfies both criteria (Figure 2j).

As a more stringent test of OPM structure, we counted the number of periodic pinwheel-like discontinuities in the OPM [41] and compared to the expected value of ~3.1 pinwheels / *mm*^2^ in macaque V1 [66]. Multiple pinwheels are apparent in both the TDANN and the Hand-Crafted SOM (Figure 2k). To facilitate quantitative comparison across models, we compute pinwheel *density* – the number of pinwheels normalized by the average spacing between “columns”, i.e. clusters of units preferring the same orientation. We find that the TDANN has lower pinwheel density (range across seeds = [2.0, 2.3] pinwheels / column spacing^2^) than macaque V1, but significantly higher than either the Task Only ([0.2, 0.8]; Mann-Whitney *U* = 25, *p* = .008) or Unoptimized models (0 pinwheels; Figure 2k). The Hand-Crafted SOM has higher pinwheel density ([3.7, 4.5]) than the TDANN, but the DNN-SOM has no detectable pinwheels. Although the TDANN has pinwheel density approaching that of macaque V1, we note that the orientation column spacing in the TDANN (~ 3.5mm width) does not match macaque V1 (~ 1mm). This mismatch, caused in part by our commitment of the TDANN as a model of human visual cortex and not macaque visual cortex, can also be overcome by increasing the number of units in the network at the expense of increased computational cost (Supplementary Figure S5).

#### The TDANN predicts maps of spatial frequency and color preference in V1

While OPMs are the best-studied feature of V1 functional organization, the cortical sheet simultaneously accommodates organized maps of spatial frequency [43] and chromatic tuning [71, 38]. An accurate model of V1 should also predict these aspects of V1 functional organization. We compared spatial frequency preference maps in macaque V1 (data from [43]) and in the TDANN V1-like layer and found a smooth progression of preferred spatial frequency in both (Figure 2f). Quantifying the difference in spatial frequency tuning as a function of cortical distance indicates that the TDANN map ([min, max] of smoothness across random initializations = [.38, .54]) is as smooth as the map in macaque V1 (0.53; Figure 2g), whereas maps from Task Only ([.23, .36]) and Unoptimized models ([.02, .03]) are far less smooth than macaque V1, and both the Hand-Crafted SOM ([.79, .81]) and the DNN-SOM ([.83, .86]) are again far smoother than the neural data. We observe similar results for maps of chromatic preference (Figure 2h, i), where comparisons are made to imaging of cytochrome oxidase (CO) uptake that is prevalent in color-tuned neurons (data from Livingstone and Hubel [38]). In the TDANN chromatic map, the fraction of units with opposite color-tuning increases with cortical distance, again exhibiting comparable smoothness to macaque V1 (TDANN smoothness: [.38, .54], macaque: .53). Together, our analyses demonstrate that the TDANN predicts the multifaceted functional organization of macaque V1, providing a stronger match to neural data than existing models such as the standard Hand-Crafted SOM.

### The TDANN reproduces the functional organization of higher visual cortex

Because benchmarks measuring the topographic similarity between models and higher visual cortex, i.e. primate inferior temporal (IT) and human ventral temporal cortex (VTC), are still underdeveloped, we introduce five quantitative benchmarks that compare both responses and topography. Response properties are compared by measuring the similarity of population category selectivity patterns with representational similarity analysis (RSA; Kriegeskorte et al. [72]), as in Margalit et al. [73], Haxby et al. [74]). Topographic properties are then compared against four complementary benchmarks: 1) the smoothness of category selectivity maps, 2) the number of category selective patches, 3) the area occupied by those patches, and 4) the spatial overlap of units selective for different categories. We compute these metrics for the TDANN’s VTC-like layer and for VTC data from eight human subjects in the Natural Scenes Dataset (NSD) [75] (Supplementary Figure S6). We also evaluate two alternative models of VTC topography: an SOM trained on the outputs of a categorization-pretrained AlexNet (DNN-SOM, cf Doshi and Konkle [13], Zhang et al. [12]) and a variant of the Interactive Topographic Network (ITN) that is trained on the same dataset (ImageNet) we used (Blauch et al. [20]: Supplementary Figure S19C). Human subjects and models were all presented a common set of 1,440 object category images [76] composed of five categories: faces, bodies, written characters, places, and objects (cars and instruments). Selectivity was computed as the *t*-value for each category, for each human voxel and model unit.

#### The TDANN predicts patterns of category selectivity

We characterize neuronal responses in VTC by computing a representational similarity matrix (RSM): the similarity between pairs of distributed selectivity patterns to each of the five object categories. The average RSM from human VTC indicates high similarity between patterns of selectivity for faces and bodies, and low similarity between selectivity for faces and places (Figure 3a). The alignment between any two RSMs is computed as Kendall’s *τ*. RSMs from different subjects and hemispheres were very similar, with the 95% CI of Kendall’s *τ* = [.72, .75]. We then compute RSMs for each model and compare against the human data, finding that some models provide a closer match to human VTC than others (ANOVA *F* (4, 331) = 630; *p* < 10^−152^). TDANN RSMs closely mirror those in human VTC (*τ* = [.69, .73]), significantly better than DNN-SOM (*τ* = [.31,.35]; post-hoc Tukey’s HSD *p* < 10^−13^), ITN (*τ* = [.46,.56]; *p* < 10^−13^), Task Only (*τ* = [.65,.68]; *p* = .001) and Unoptimized (*τ* = [.11,.14]; *p* < 10^−13^) models (Figure 3b). The similarity between human and TDANN RSMs also depends strongly on the training data being naturalistic. Training on artificial stimuli such as white noise and sine gratings yields RSMs that significantly deviate from the human data (Supplementary Figure S9b).

**Figure 3.**
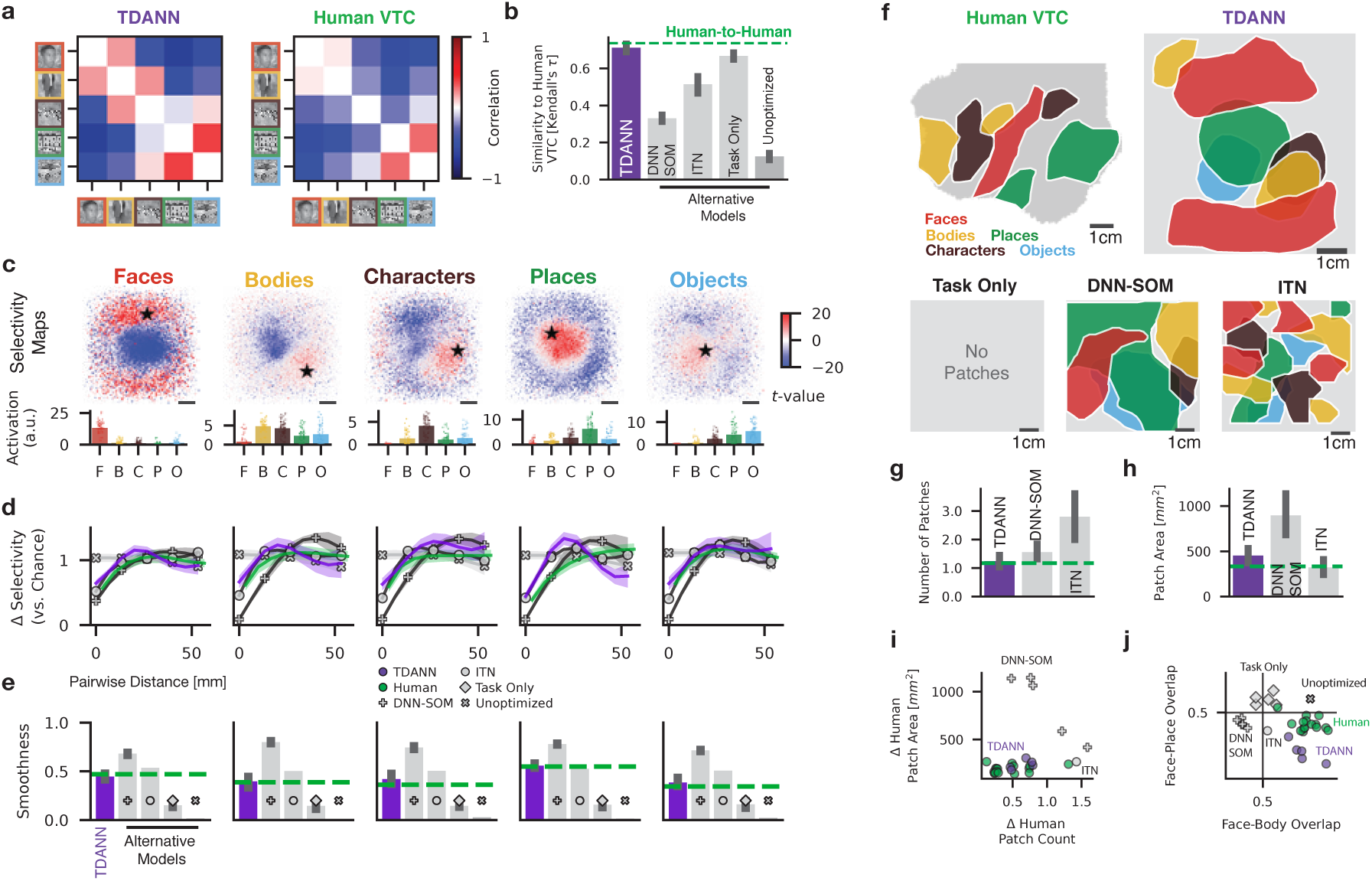
The TDANN predicts the functional organization of higher visual cortex. **(a)** Representational similarity matrices (RSMs) for the TDANN and human VTC, computed across selectivity maps of the five object categories. Diagonal is blank to indicate trivially perfect correlation. **(b)** Functional similarity between the TDANN, human VTC, and alternative models, measured as the similarity of RSMs. Green: mean of pairwise human-to-human similarity values. **(c)** Selectivity (*t*-value), for each category plotted on the simulated cortical sheet of the VTC-like layer in an example TDANN. Black star: unit whose responses to images in each of the five categories are plotted directly below (individual dots: single images, bar height: mean across images). Scale bar: 1cm. **(d)** Difference in pairwise selectivity as a function of pairwise cortical distance for units in each of five candidate model types: the TDANN (purple), deep neural network self-organizing map (DNN-SOM; plus markers), interactive topographic network (“ITN”, Blauch et al. [20]; circles), Unoptimized (“x” markers), and Task Only (diamond markers). Curves are normalized to the chance level obtained by random sampling of unit pairs. Green: Human data averaged over the eight subjects in the NSD data. Shaded regions: 95% confidence interval across different subsets of units from models trained with different random initial seeds. **(e)** Smoothness of selectivity maps for each category and each candidate model. Dashed green: mean of human data. **(f)** Category-selective patches for an example hemisphere in human ventral temporal cortex (VTC), TDANN, a Task Only model (no patches detected), a DNN-SOM, and a reproduction of the “ITN” simulated cortical sheet from [20]. Object categories are indexed by color as in (a) and (c). Examples from different initial random seeds are shown in Supplementary Figure S10. **(g)** Number of category-selective patches (averaged across categories) for the TDANN, DNN-SOM, and ITN. Dashed green: average of human data. ANOVA for difference in patch count: *F* (5, 179) = 32.7, *p* < 10^−22^. Post-hoc Tukey’s tests: significant difference between VTC and ITN (*p* = 1.2 *×* 10^−5^). **(h)** Average surface area of category-selective patches. Same plotting conventions as in (f). ANOVA for difference in patch area: *F* (5, 187) = 15.4, *p* < 10^−11^. Post-hoc Tukey’s tests: significant difference between VTC and DNN-SOM (*p* < 10^−10^). **(i)** Each human subject and model instance compared to the mean patch area (y-axis) and patch number (x-axis) in the human data. **(j)** Overlap between face-selectivity and body-selectivity vs. overlap between face-selectivity and place-selectivity, for each human hemisphere (green dots), each TDANN instance (purple dots), the ITN (gray dot), each DNN-SOM (gray plus signs), and Task Only models (gray diamonds).

#### The TDANN predicts category-selectivity maps

To compare models against topographic benchmarks, we generate selectivity maps for each of the five object categories (Figure 3c), then quantify their structure by measuring the pairwise difference in selectivity as a function of pairwise cortical distance (Figure 3d). We find that for all categories, the curve computed for TDANN is similar to human VTC, whereas the DNN-SOM and ITN are abiologically smooth, and maps in the Unoptimized and Task Only models lack structure. We summarize category selectivity map structure with the same smoothness metric used in V1 (Figure 3e), and find that TDANN maps were as smooth as those in human VTC (permutation test: *p* = .30). In contrast, VTC maps were significantly smoother than Task Only or Unoptimized models (*ps* < .001) and less smooth than the DNN-SOM (*p* < .001). ITN category selectivity maps were smoother on average than VTC, but not significantly so (*p* = .10).

For the remaining topographic benchmarks, we follow the literature by thresholding selectivity maps to find strongly-selective units (Supplementary Figure S6a-d). Clusters of selective units are identifiable in human VTC, TDANN, the SOM and ITN models, but not in Task Only or Unoptimized models. We use a data-driven approach to automatically identify large contiguous clusters of selective units as “patches” (Figure 3f). We find similar sets of patches in VTC and the TDANN: both contain a small number of patches selective for each category (except for object-selective patches, which are not found in VTC), and the patches are similar in size. Quantitative comparison supports the similarity of human VTC and TDANN: there is no significant difference in patch count (*p* = 0.99, Figure 3g) or patch area (*p* = 0.67; Figure 3h). In contrast, we find that the ITN has more than twice as many patches as VTC (*p* = 1.2 *×* 10^−5^), although the patches are as large on average as those in VTC (*p* = 0.99). The DNN-SOM fails to match VTC in the other extreme: while the number of patches in the DNN-SOM is similar to that in VTC (*p* = 0.15), the patches are too large (*p* < 10^−10^). Joint comparison of models and humans on both patch count and size (Figure 3i) highlights the stronger correspondence between TDANN and human VTC than alternative models.

An important hallmark of the functional organization of higher visual cortex is the reproducible spatial arrangement of units selective for different categories. A prominent example is the close proximity of face-selective and body-selective regions [49, 77] and the separation between face- and place-selective regions. A measure of proximity between face- and body-selective regions was previously introduced in Lee et al. [78]. Here we measured the co-occurrence of face-selective and body-selective units (and face-selective and place-selective units) in human VTC with an overlap score that ranges between 1 (face-selectivity perfectly predicts body-selectivity) to 0.5 (no relationship), to 0 (face- and body-selectivity perfectly anti-correlated). As expected, Face-Body overlap scores are high in human VTC (95% CI across subjects and hemispheres: [.66, .72]), whereas Face-Place overlap was significantly lower (95% CI: [.40, .45], Wilcoxon signed-rank test against one-sided alternative *W* = 136; *p* = 1.5 *×* 10^−5^; Figure 3j). The same pattern is apparent in the TDANN: Face-Body Overlap ([.63, .71]) is significantly higher than Face-Place Overlap ([.14, .26]; *W* = 15; *p* = .03). In the ITN, the Face-Body overlap score was lower than in human VTC (.52), but still higher than the Face-Place overlap score (.36). Neither the the DNN-SOM nor the Task Only models had higher Face-Body overlap than Face-Place overlap (Figure 3j; *ps* > 0.5).

To further gain intuition for the tuning profiles of model units, we synthesized images that optimally drive each region of the VTC-like layer. We find that the VTC-like layer smoothly maps object feature space onto the two-dimensional simulated cortical sheet; e.g., face-patches are optimally driven by stimuli with apparent eyes (Supplementary Figure S7). We also tested how the nature of the training dataset affects the accuracy of topographic maps in the TDANN (see Lee et al. [78], Figure 7 for a similar analysis). We find that training the TDANN on natural images (either ImageNet [64] or Ecoset [79]) produces accurate V1-like and VTC-like maps, whereas training on noise or simpler hand-crafted stimuli fails to provide a unified account of ventral stream topography (Supplementary Figure S9).

Together, these results demonstrate that TDANN is the only model to exhibit spatially structured category selectivity that is consistent with a large battery of benchmarks comparing models to human VTC.

### Multiple signatures of functional organization emerge at the same spatial constraint strength

The TDANN optimization framework requires the selection of a single free parameter, *α*, the weight of the spatial loss in the training objective. When *α* = 0 (“Task Only”), spatial information is ignored during training, whereas setting *α* too high may encourage pathologically strong correlations that interfere with representation learning. In the results above, *α* is set to 0.25. Here, we validate this choice by demonstrating that many benchmarks of neural similarity are simultaneously satisfied by low-to-intermediate values of *α*.

Comparison of OPMs in the V1-like layer and category-selectivity maps in the VTC-like layer (Figure 4a) in models trained at 7 different levels of *α* shows that functional organization is absent when *α* = 0, structured at intermediate values of *α*, and deteriorates at the highest values of *α*. We quantify the dependence of functional organization on *α* with three kinds of benchmarks: functional similarity (Figure 4b), map smoothness (Figure 4c), and presence of topographic phenomena (i.e. pinwheels and patches; Figure 4d). First considering functional similarity, we find that the fraction of V1-like layer units that are orientation selective is closest to macaque V1 when *α* is low, and representational similarity between the VTC-like layer and human VTC is maximized at *α* = 0.25 (Figure 4b). The smoothness of topographic maps is most brain-like at *α* = 0.1 for OPMs in the V1-like layer and at *α* = 0.25 for category-selectivity maps in the VTC-like layer (Figure 4c). Finally, we find that the density of pinwheels in the V1-like layer and category-selectivity maps in the VTC-like layer are most similar to measurements in macaque V1 and human VTC, respectively, at *α* = 0.25 (Figure 4d).

**Figure 4.**
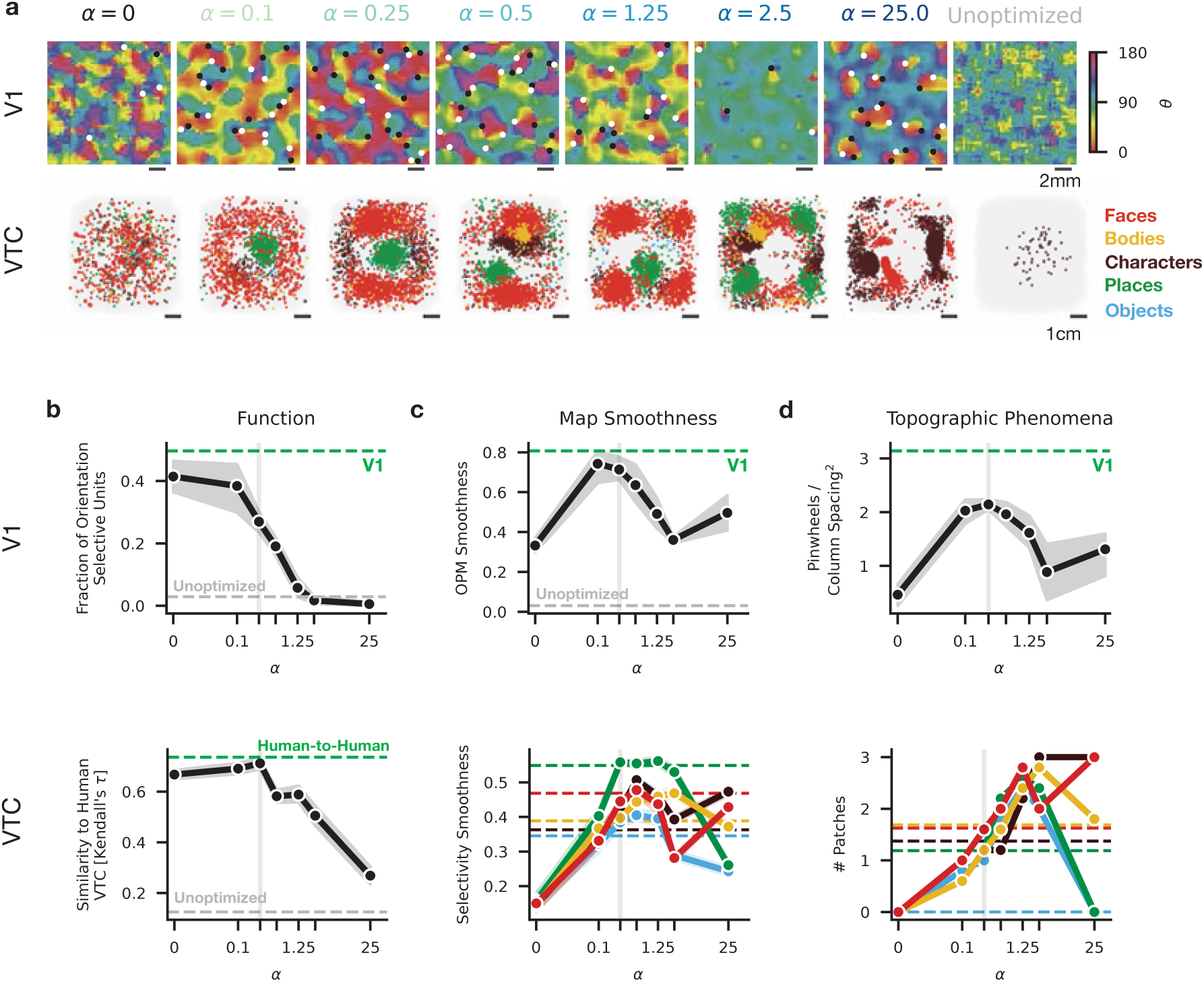
Convergence of multiple benchmarks indicates a balancing between functional and spatial constraints. **(a)** Topographic maps in the V1-like (top row) and VTC-like layer (bottom row) of TDANN models trained at different levels of the spatial weight *α*. Top: Orientation map structure and pinwheels become apparent at *α >* 0.1 and persist until *α* = 1.25. Dots: estimated pinwheel locations; black: clockwise, white: counterclockwise. Bottom: Category selectivity maps, with selective units (*t >* 12) colored according to their preferred category. **(b)** Functional correspondence to neural data as a function of *α*. Top: Fraction of units strongly orientation selective (circular variance *≤* 0.6) in the V1-like layer. Dashed green: value measured in macaque V1 (from Ringach et al. [34]). Dashed gray: mean value for Unoptimized models. Shaded regions: 95% CI across multiple initial random seeds. Bottom: Representational similarity between the VTC-like layer and human VTC (as in Figure 3). Error region indicates 95% CI across model seeds and human hemispheres. In both plots, the vertical line at *α* = 0.25 marks the default value used in prior figures. **(c)** Topographic map smoothness as a function of *α*. Top: OPM smoothness in the V1-like layer. Dashed green: value in macaque V1. Dashed gray: smoothness in an Unoptimized model. Bottom: Category selectivity map smoothness in the VTC-like layer. Dashed lines indicate means across human subjects and hemispheres from the NSD data; one line per category. **(d)** Density of topographic phenomena of interest as a function of *α*. Top: Pinwheel density in OPMs from the V1-like layer, as a function of *α*. Bottom: Number of category selective patches for each category in the VTC-like layer, as a function of *α*. Human data in dashed lines.

A specific range of *α* values (0.1 *≤ α ≤* 0.25) thus produces experimentally-observed outcomes across a variety of independent functional and topographic benchmarks in multiple brain areas, suggesting that the *α* parameter may provide insights into biophysical mechanisms underlying the emergence of functional organization.

### Two key factors underlying functional organization: self-supervised learning and a scalable spatial constraint

Having established that specific TDANN models accurately predict the functional organization of the ventral visual stream, we consider what key factors enable the emergence of this functional organization. We reasoned that if some combinations of optimization objectives yield brain-like functional organization and others do not, it will shed light on the constraints underlying the observed functional organization. Thus, we train models with alternative task and spatial objectives, then apply our benchmarks to evaluate which models are most consistent with empirical data.

For the “task component” of its loss function, the TDANN uses contrastive self-supervision [61, 63], a framework for learning representations that transfer easily to many downstream tasks. These self-supervised algorithms have been shown to generalize to many downstream computer vision tasks despite being trained only on a large set of unlabeled natural images [80]. However, most studies comparing neural networks to the brain have used a supervised object categorization ([26, 25, 78]; Figure 5a-bottom left). Thus, we tested whether training with an object categorization objective produces different functional organization than self-supervision, and if so, which is more similar to the observed functional organization of the ventral visual stream.

**Figure 5.**
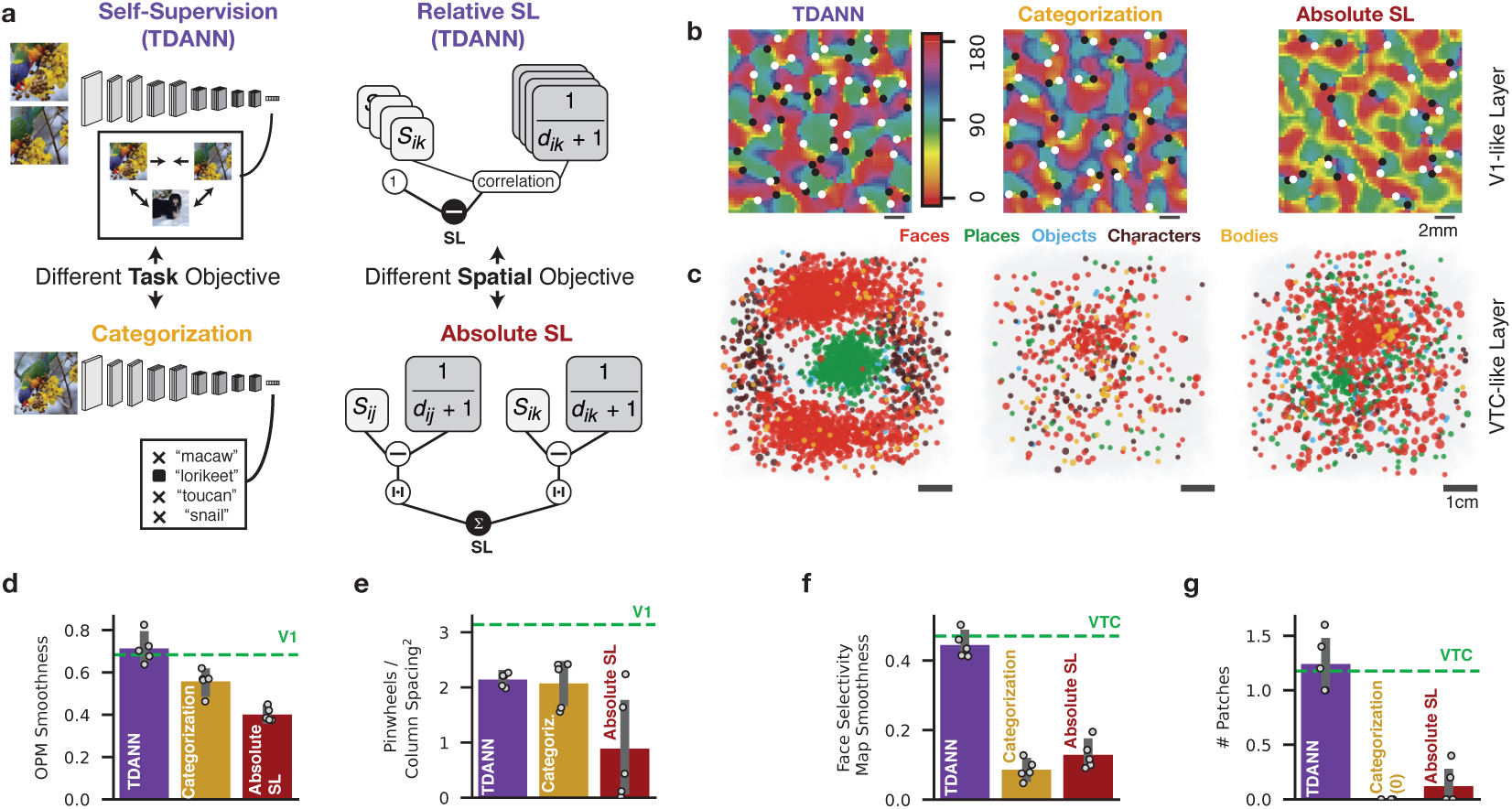
Self-supervision and scalable spatial constraints underly the emergence of functional organization. In each panel, TDANN shown in purple, Categorization-trained in gold, Absolute SL in red, and ventral stream measurements in green. **(a)** Left: comparison of task objectives. The TDANN uses contrastive self-supervision (top) which encourages similarity between representations of different views of the same image while increasing distance between representations of views of other images. Categorization (bottom) compares predicted class probabilities to the human-labeled correct class. Right: comparison of spatial objectives. *S_ij_* : response similarity of units *i* and *j*. *d_ij_* : cortical distance between units *i* and *j*. TDANN uses the Relative SL (top), which correlates the population of response similarities and pairwise inverse distances. Prior work [78] used the Absolute SL (bottom), which directly subtracts inverse cortical distance from response similarity magnitude. **(b)** Smoothed orientation preference maps (OPMs) in the V1-like layer of the TDANN (left), a Categorization trained model (middle), and a model trained with the Absolute SL (right). Dots: detected pinwheels. *α* = 0.25 for models shown in each panel. **(c)** Category selective units in the VTC-like layer of the TDANN (left), a categorization trained model (middle) and a model trained with the absolute SL (right). **(d)** Right: Smoothness of OPMS in the V1-like layer of each model type. Green line: value computed macaque V1. **(e)** Density of detected pinwheels. Green: estimated value in macaque V1. **(f)** Right: Smoothness of face selectivity maps in the VTC-like layer of each model type. Green line: value from human VTC. **(g)** Average number of category-selective patches, in the VTC-like layer in each model. Green: average value in human VTC.

We also investigate how the form of the spatial objective function affects emergent functional organization. The spatial component of the TDANN loss function is generally intended to capture the constraints on unit-to-unit correlations within cortical neighborhoods, but the specifics of its functional form embody conceptually distinct mechanistic ideas about how a hypothetical cortical development circuit might measure functional correlations and compare them to cortical distances. In prior work, Lee et al. [78] introduced a spatial loss function that subtracts the inverse of pairwise cortical distances from the magnitude of pairwise response correlations (Figure 5a-bottom right), such that nearby units develop similar responses. That loss function was developed to match empirical measurements in macaque IT, but was not intended to generalize to other regions of the human ventral visual stream. We refer to it as the Absolute Spatial Loss (or *SL*_Abs_), because minimizing it requires an absolute match between response correlations and the inverse of cortical distances. While Lee et al. [78] found that training models with *SL*_Abs_ produced clustering of category-selective units in a late model layer, we discovered a critical flaw when training with *SL*_Abs_ in all model layers: in layers with shorter cortical distances, *SL*_Abs_ can only be minimized if response correlations are pathologically high. The TDANN instead uses a more flexible spatial loss function that we term the Relative Spatial Loss (*SL*_Rel_; Figure 5a-top right). This SL requires that inverse cortical distances will be correlated with response similarity (see Methods for mathematical details). *SL*_Rel_ effectively enforces response similarity between pairs of units that are *relatively* close together. Thus, the Relative SL allows the distance over which local correlations extend to depend on the total size of the cortical area. Interestingly, we find that switching from *SL*_Abs_ to *SL*_Rel_ slightly increased the model’s capacity for object categorization at all levels of *α* (Supplementary Figure S12). How do models trained for different objectives differ on topographic benchmarks?

We compare the TDANN (self-supervised and Relative SL) to categorization-trained models (differing only in task objective) and Absolute SL models (differing only in spatial objective) on our battery of topographic and functional benchmarks: (i) evaluating the smoothness of OPMs and face-selectivity maps in the V1-like and VTC-like layers, respectively, and (ii) counting the number of pinwheel-like discontinuities and category-selective patches in those layers, respectively. Categorization-trained models were slightly but significantly less smooth than the TDANN (mean smoothness = 0.56, *U* = 25, *p* = 0.008), but with an equal density of pinwheels (2.07 pinwheels / column spacing ^2^; *U* = 10, *p* = 0.69). Absolute SL models generally resemble those in the TDANN (Figure 5b), but with significantly lower smoothness (TDANN mean: 0.71, Absolute SL: 0.40; *U* = 25, *p* = 0.008; Figure 5d) and slightly lower pinwheel density (TDANN: 2.14 pinwheels / column spacing ^2^, Absolute SL: 0.89; *U* = 21, *p* = 0.09; Figure 5e).

Strikingly, however, category-selectivity maps in the VTC-like layer were much less organized in the Categorization-trained models than in the self-supervised TDANNs. At the same spatial weight of *α* = 0.25, clear clusters of category-selective units are observed in the self-supervised but not the categorization-trained model (Figure 5c). The Absolute SL models also fail to form organized category-selectivity maps at this level of *α*. Quantitative comparison reveals smoother category selectivity maps in the TDANN (mean smoothness of face-selectivity maps = 0.44) than in either categorization-trained models (0.09; Mann-Whitney *U* = 25, *p* = 0.008; Figure 5f) or in Absolute SL models (0.13). The TDANN also has a significantly higher number of identified category selective patches (mean = 1.2) than either categorization-trained (mean = 0) or Absolute SL alternatives (mean = 0.08; *U* = 25, *p* = 0.008; Figure 5g). Thus, the nature of the training objective strongly constrains the emergent functional organization, with self-supervised learning and relative spatial loss objectives producing the most brain-like functional organization.

### Spatial constraints make learned representations more brain-like by reducing intrinsic dimensionality

A natural question is whether training for spatial objectives also has an effect on the *non-topographic* properties of learned representations. Because the TDANN allows the network’s features to be influenced by the spatial constraint during training, we can directly address this question.

A powerful way to test if spatially-constrained models learn different features than standard DANNs is to measure how well model unit responses can predict neural responses to large set of naturalistic images in primate visual cortex [30, 28, 25, 81]. A popular approach to predicting neuronal firing rates is to fit the responses of individual neural units with a linear combination of many hundreds or thousands of model units. Consistent with prior work involving non-spatial models [61], we find that models trained with different objectives are largely indistinguishable in their ability to predict neural firing rates when using this standard linear-regression method for mapping model units to neural firing rates [25, 61, 29] (Figure 6a). The linear-regression mapping is thus insensitive to the dramatic differences between models trained with different objectives and spatial constraint magnitudes that are apparent in our analysis of functional organization. A possible explanation for this apparent discrepancy is that linear regression is too permissive of mapping: even if a model lacks individual units that resemble recorded neurons, a combination of units might still allow for accurate prediction of neural responses. We tested this prediction by performing a more stringent one-to-one mapping, in which individual VTC-like layer model units – not a linear mixture of units – are assigned to individual VTC voxels in a one-to-one fashion. Intriguingly, we found that this one-to-one assignment resulted in much stronger matches between TDANN model units and voxels recorded in the Natural Scenes Dataset (NSD) [75] than models trained with other objectives (i.e. categorization or Absolute SL, Figure 6b). This correlation peaks at *α* = 0.25, the same value identified by topographic benchmarks (Figure 4), providing more evidence that the constraints driving brain-like functional organization also make learned representations more brain-like.

**Figure 6.**
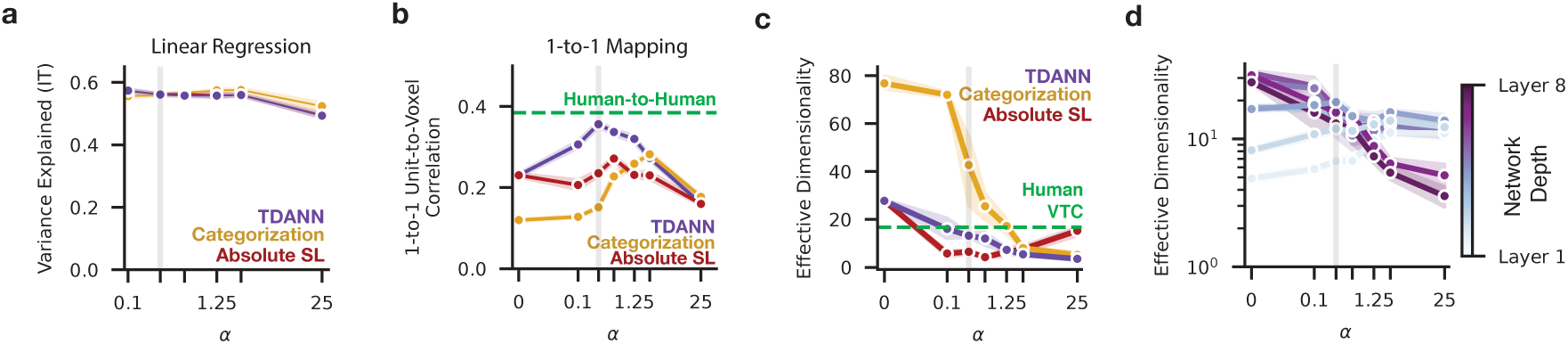
Spatial constraints make learned representations more brain-like and reduce intrinsic dimensionality. **(a)** Variance explained under a linear regression mapping between model units and macaque IT neurons, as a function of the spatial loss weight *α* and the training objective. **(b)** Mean correlation between model units and VTC voxels under a one-to-one mapping as a function of *α*. Green: mean human-to-human correlation under the same one-to-one mapping. **(c)** Estimated effective dimensionality (cf. Elmoznino and Bonner [83], Del Giudice [84]) of the population response in the VTC-like layer of models trained at different levels of *α* and with different objectives. Green: mean value in human VTC from the NSD dataset. **(d)** Effective dimensionality in the TDANN across all layers and levels of *α*. In all panels, shaded vertical bar indicates value of *α* demonstrated in prior analyses to best match topographic phenomena.

Many factors might contribute to the differences in representation between the TDANN and those of poorer-fitting models. Because the TDANN’s spatial constraint encourages units to respond more similarly to one another, we hypothesized that the intrinsic dimensionality of the population might decrease as *α* increases. Relatedly, recent work has demonstrated that spatially unconstrained DANN responses to natural images have substantially higher intrinsic dimension than real macaque and rodent V1, and that models with lower dimensionality better predict neural responses [82]. Thus, we tested whether decreased intrinsic dimensionality might explain why the TDANN representations are more brain-like than representations from other models. Consistent with our hypothesis, we find that the addition of the spatial constraint decreases intrinsic dimensionality in the VTC-like layer regardless of the training objective (Figure 6c; see Supplementary Figure S13a for eigenspectra in all layers). When *α* = 0, all models have higher effective dimensionality (ED; Elmoznino and Bonner [83], Del Giudice [84]; see methods) than human VTC (mean across subjects = 16.7), although the dimensionality of the VTC-like layer in categorization-trained models (76.8) is nearly three times higher than in the self-supervised models (TDANN and Absolute SL: 27.8). At the spatial weight magnitude *α* = 0.25, at which the TDANN best matches neural data, the TDANN’s VTC-like layer approaches the dimensionality of human VTC (TDANN mean = 13.2). However, the dimensionality of models trained with *SL*_Abs_ decreases too quickly (mean = 6.5), and categorization-trained models remain higher than human VTC at this level of *α* (mean = 42.7).

We conclude that the close match between the TDANN and human VTC, on both topographic and non-topographic benchmarks, may be due in part to an alignment of their intrinsic dimensionality. Similar results are observed when summarizing the response eigenspectrum with power law fits, as in Stringer et al. [85], Kong et al. [82] (Supplementary Figure S13c). Intriguingly, we find that the effective dimensionality of the TDANN roughly converges to a common value of approximately 15 across model layers at *α* = 0.25 (Figure 6d), raising the possibility that a similar dimension stabilization phenomenon occurs across brain areas in the ventral stream. These results provide new evidence that the computational constraints generating cortical topography strongly influence *non-topographic features*, making them more brain-like by virtue of decreasing the dimensionality of population responses.

### The TDANN minimizes inter-layer wiring length

Identifying the optimization paradigm that is most consistent with neural data provides insight into the constraints underlying neural development, but prompts a deeper question: why would these constraints be favored by evolutionary selection? A natural hypothesis is that cortical networks with strong functional organization also minimize wiring length, and thus reduce brain size, weight, and power consumption [86, 18]. We test this hypothesis by asking whether the optimization paradigm that generated a functional organization that best fit neural benchmarks – intermediate spatial weight *α*, self-supervised learning, and spatial costs that scale with cortical surface area – also reduces between-layer wiring length. In feedforward networks that lack intra-layer connectivity, such as the TDANN, any gains in wiring efficiency must be between layers. Accordingly, we measure inter-layer wiring length by identifying populations of co-activated units in adjacent layers, then estimating the length of fibers needed to connect those populations. We first present natural images to the network and record the locations of the most responsive units in each layer, then simulate fiber bundles that originate in an earlier “source” layer and terminate in the following “target” layer, adding inter-layer fibers until the total squared distance between each activated unit and its nearest fiber is below a specified threshold (see Methods, Figure 7a). The total wiring length is taken as the sum of the lengths of each fiber.

**Figure 7.**
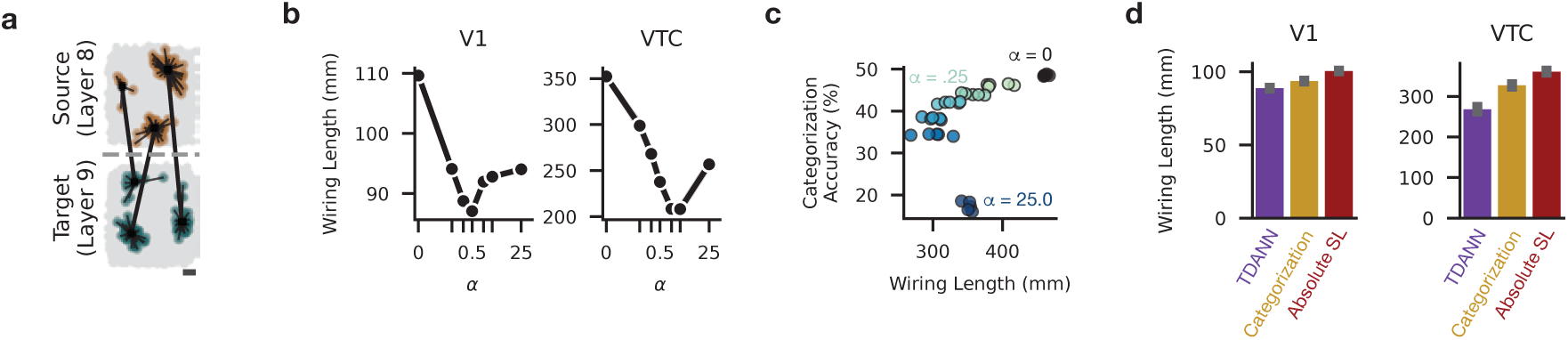
Minimization of inter-layer (feedforward) wiring length in models with brain-like functional organization. **(a)** Example wiring length computation between adjacent layers. Units in brown are the top 5% most active units in the Source layer for an arbitrarily-selected natural image, while units in green are the top 5% most active in the Target layer. Black dots show the origination and termination points of fibers that would be required to connect populations of active units across layers. **(b)** Wiring length between layers 4 and 5 (“V1”; left), and layer 8 and 9 (“VTC”, right) as a function of *α*. Shaded regions: 95% CI of measurements from different cortical neighborhoods, model seeds, and input images. **(c)** Accuracy on object categorization vs total wiring length, for models trained at different levels of *α*. **(d)** Wiring length in both early and later model layers for models trained with different task and spatial objectives (*α* = 0.25 for all). Error bar: 95% CI over different image presentations and model seeds.

Presenting the TDANN with natural images leads to clustered responses in the VTC-like layer of all models trained with *α* > 0, with multiple clusters apparent at higher levels of *α* (Supplementary Figure S14). Does the increase in clustering within layers result in shorter wiring length between layers? We find that inter-layer wiring length is indeed minimized at higher levels of *α* (Figure 7b). However, we also find that object categorization performance decreases as wiring efficiency improves (Figure 7c), indicating that models at low-to-intermediate levels of *α* optimally balance performance with inter-layer wiring efficiency. This coincidence of optimal *α* values suggests that the functional organization of the ventral visual stream balances inter-area wiring costs with performance. Critically, we find that wiring is most efficient for the optimization objectives that yield the most brain-like functional organization: wiring length is higher in both categorization-trained models and those trained with the Absolute SL (Figure 7d).

Thus, wiring length minimization provides a normative explanation for the superiority of self-supervised learning and area-normalized spatial constraints.

### Proof-of-principle: Using the TDANN as a digital twin for experimental design

A quantitatively accurate and mechanistically grounded model of functional organization, such as the TDANN, enables a spectrum of applied use cases that rely on estimating the effects of spatially-modulated neural perturbations. Here we apply TDANN as a digital twin of visual cortex and demonstrate two novel applications: 1) performing an *in silico* microstimulation experiment, and 2) proof-of-principle for prototyping a simple cortical prosthetic device.

#### Simulated microstimulation reveals functional similarity of conneted unit populations

Microstimulation experiments in the macaque [87] found that stimulating neurons in a face patch selectively drives activity in other face patches, and prior work with topographic models of macaque IT [78] found a similar result. We tested if the TDANN also captures this connectivity by stimulating local populations of units in the penultimate model layer and recording evoked responses in the following VTC-like layer. Mirroring results in macaque IT, we find that stimulating units in a TDANN face patch drives localized activity in a face patch in the following layer (Figure 8a). We repeat the stimulation for 99 other sites equally spaced on the simulated cortex, and find that the selectivity of a stimulated unit in the source layer strongly predicts the selectivity of activated units in the target layer (Figure 8b), especially for stimulation sites closer to the center of the simulated cortical tissue.

**Figure 8.**
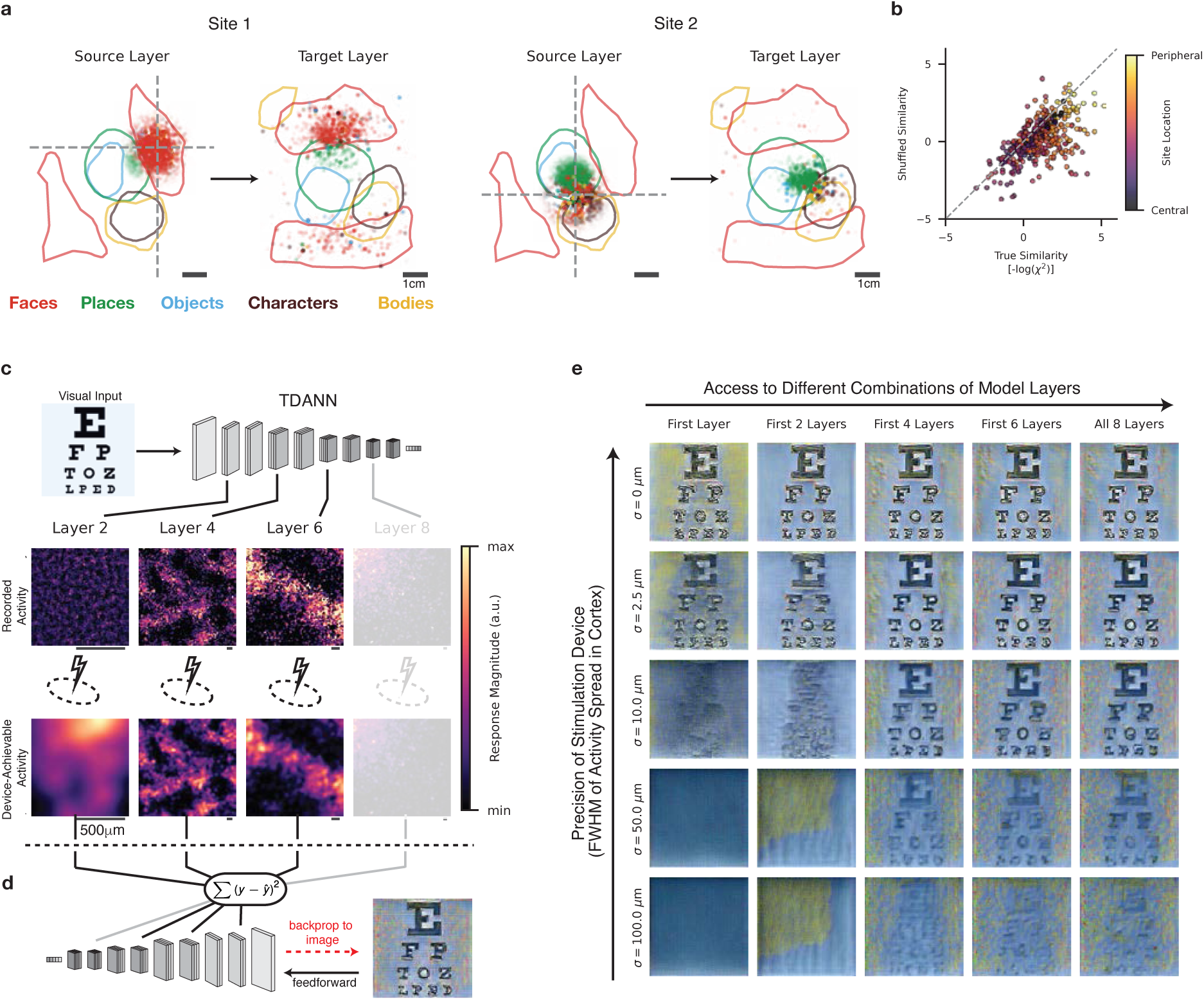
Using TDANNs to simulate spatial stimulation devices. **(a)** Stimulation of a local population of units in the second to last convolutional layer drives spatially-localized responses in the final convolutional layer. Responses are functionally aligned, such that stimulating face-selective units (Site 1) drives activity in face-selective units in the following layer. Right: Results for a second stimulation site, at the intersection of place-, body-, and character-selective patches. **(b)** Similarity in tuning of stimulated units in the source layer and responding units in the target layer for 100 evenly-spaced stimulation sites. Each dot compares tuning similarity for the true distribution of activated units (x-axis) and a randomly shuffled selection of units (y-axis). Dot color: distance of the stimulation site from the center of the cortical tissue. **(c-d)** Conceptual framework for applying the TDANN to the prototyping of visual cortical prostheses. **(c)** Stimulation Simulator: the TDANN is used to generate predicted activity patterns from a given visual input (top row). Patterns are then degraded according to the limitations on a hypothetical stimulation device: reduced spatial precision results in blurring of the target activity pattern (bottom row), and limits to regional access restrict the set of layers that participate. Here, Layer 8 is faded-out to show that this particular hypothetical device cannot reach that cortical area. **(d)** Given a device-achievable stimulation pattern produced by the Stimulation Simulator in (c), we synthesize the image that could evoke that pattern: the predicted percept. To build intuition for the fidelity of predicted percepts, we use an example input image of the the first four lines of a Snellen eye chart. **(e)** Predicted percepts for 25 theoretical cortical stimulation devices with different capabilities. Devices vary in the precision with which they are able to produce desired activity patterns (full-width at half-maximum (FWHM) of the spread of activity on cortex increases with rows) and the number of cortical areas that can be simultaneously simulated (columns).

#### Simulation of cortical prosthetic devices with TDANNs

A unique advantage of a unified topographic model such as TDANN is that it can be used to prototype the effects of simultaneous stimulation of multiple cortical areas, experiments which are challenging to perform *in vivo*. Based on recent advances in machine learning and visual cortical prostheses [88, 89, 90], we introduce a framework using TDANNs to prototype multi-region cortical stimulation devices. The framework has two components (Figure 8c, d): 1) a Stimulation Simulator that transforms desired activity patterns on the cortical sheet into *device-achievable* patterns, and 2) a Percept Synthesizer that estimates the percept evoked by stimulation with those patterns.

The Stimulation Simulator takes an input image, uses the TDANN to predict the precise pattern of responses in each layer, and then constrains that pattern into one that is physically achievable by a specific hypothetical stimulation device (Figure 8c). We model two kinds of physical constraints: spatial precision – the resolution at which the device can create activity patterns, and regional access – the subset of cortical areas that are accessible to the device. Spatial precision is modeled as a Gaussian blur of the desired activity pattern and regional access by restricting the model layers that participate in the simulation.

To synthesize percepts from device achievable patterns, we use an approach inspired by Granley et al. [90] and Shahbazi et al. [91] to synthesize the input image which generates the target activity pattern – i.e., a neural metamer. Figure 8e illustrates predicted percepts for hypothetical cortical stimulation devices with variable precision and access. Unsurprisingly, a device with infinitely high spatial stimulation precision yields sharp percepts even when only early cortical areas are stimulated (Figure 8e, top left). However, the percepts quickly deteriorate as the spatial precision of the device decreases (Figure 8e lower left). Notably, our simulation suggests that, at lower spatial precision, the quality of percepts can be improved by adding stimulation of higher cortical areas (Figure 8e, middle rows).

While we have neglected many critical details here, including spatiotemporal processing, cortical magnification, and the need to validate percepts, we hope that this proof of principle motivates the use of TDANN to make testable predictions about the nature of percepts elicited by various cortical stimulation devices.

## Discussion

In this work, we leveraged the neural network modeling framework to seek the principles of functional organization in the primate ventral visual stream. We found that training a spatially-augmented deep neural network for a specific combination of objectives results in a model, the TDANN, that captures topographic properties throughout the ventral stream, from the pinwheels of V1 to the category-selective patches of higher-level visual cortex.

We identified two specific factors critical to the emergence of brain-like functional organization. First, we found that self-supervised learning of task-general representations yields better organization than the more common alternative of supervising on the singular task of visual object recognition. Recent work has suggested that functional specialization in the brain – e.g., one population of units responsible for discrimination of different faces and another for recognition of different objects – arises under joint training for two different supervised recognition tasks, one for faces and one for objects [92]. Our results demonstrate that functional specialization can emerge under a single unsupervised learning objective on a single training set, suggesting that general mechanisms can produce the kinds of functional specialization that is typically assumed to require multiple objectives or multiple distinct datasets. Second, we found that the spatial constraint in our model should compare response similarity and physical similarity according to a metric that scales with the size of each cortical area, rather than being fixed for all cortical areas. This finding suggests that the actual circuits responsible for shaping the structure of local response correlation in cortical neighborhoods should scale with the surface area of each cortical region. Our identification of these two critical factors demonstrates that a goal-driven modeling approach to understanding neural sensory systems can yield concrete and specific insights into their underlying principles.

Critically, the two factors that we found are essential for brain-like functional organization in the visual system are not specific to the visual modality, and might extend to predict the abundant, yet largely unexplained, functional organization in other sensory systems. For example, neurons in primary auditory cortex are arranged according to the frequency they respond most strongly to (tonotopy [2]), and in secondary auditory areas, neurons cluster according to their preference for speech and music [23, 93]. It is possible that the representations carried by these neurons are also learned by contrastive self-supervision, and that their topographic organization is explained by scalable spatial constraints of the forms described here. Likewise, the functional organization of somatosensory [4], entorhinal [6, 5] and parietal cortices [3] may be explained by the specific yet general principles for representation learning and spatial smoothness that we have identified. Under this hypothesis, it is only the structure of the input data (e.g., auditory experience, somatosensory input) that changes, but the cortical mechanisms for learning and organization remain universal across cortical systems. Future work can directly test that hypothesis by training TDANN variants to learn spatially-organized representations specific to each system.

The TDANN is the first model to predict functional organization in multiple cortical areas by learning features and topography, from scratch, in and end-to-end optimization framework trained directly on image inputs. As such, it represents an improvement over a number of related prior approaches. For example, hand-crafted self-organizing maps (SOMs) [94, 8, 10, 11, 9] have simplified the problem of topographic map formation by modeling a limited set of fixed feature dimensions (e.g., orientation preference and spatial frequency tuning), then modifying the tuning of model units along these dimensions such that nearby units develop similar selectivity. While such SOMs produce qualitatively smooth V1-like orientation maps, we find that they fail to quantitatively predict the topographic properties of V1 orientation maps (Figure 2). Recent attempts to abandon hand-crafted feature dimensions have trained SOMs to smoothly map the outputs of categorization-pretrained DCNNs [12, 13]. While these DNN-SOMs have the advantage of operating on images rather than predefined features, we find that they are quantitatively less accurate than the TDANN (Figure 3) at explaining the functional organization of VTC, and fail to reproduce the topography of V1 (Figure 2). Another recent approach, the ITN [20], appended topographic layers to a pretrained DCNN backbone and trained for supervised categorization under an additional wiring length minimization constraint. While the ITN reproduces many features of VTC topography, it does not predict the size, number, and geometry of category-selective patches as accurately as the TDANN, and cannot predict the functional organization of areas outside VTC. Prior work from our groups also followed the TDANN optimization framework, but used a supervised categorization task, a spatial constraint that did not scale with cortical area, and applied only to the VTC-like layer [78]. While this model was able to predict many properties of the functional organization of macaque IT, it incapable of predicting the organization of other ventral stream regions. Our present results (Figure 5) demonstrate that different spatial and task objectives are required for a TDANN to accurately match the functional organization of multiple areas of the ventral visual stream.

That the TDANN is trained end-to-end provides two interesting opportunities for understanding the interaction between learned representations and functional organization during development. First, our preliminary analyses suggest that trajectories of TDANN functional architecture throughout training roughly match the faster development of earlier vs higher cortical regions (Figure S17) and the emergence of V1-like topography from retinal wave-like stimuli (Figure S18). Rigorously testing those predictions would be most interesting when the TDANN is optimized using naturalistic movie streams that match the visual statistics and acuity limitations of human development [95, 96]. Second, we found that the presence of the spatial constraint during training modulated the nature of learned representations, making them more brain-like and stabilizing their intrinsic dimensionality (Figure 6).

While the TDANN is the first unified model of ventral stream functional organization, it has a number of important limitations. Because the core DCNN architecture used in this work is strictly feedforward, there are no direct connections between different units in the same layer. Thus, we are only able to draw inferences about how the spatial constraint affects wiring length between layers. A more complex architecture could include both intra-layer recurrence and long-range feedback connections [97], although our results demonstrate that explicitly modeling these recurrent connections is not necessary to produce accurate topographic maps (see Figure 6 of Blauch et al. [20]), raising the possibility that minimization of the length of long-range fibers may be the key determinant of the functional organization of visual cortex.

We also note that our model, like all convolutional neural networks, uses the same filter weights across the entire visual field (termed “weight sharing”). This short-cut makes large-scale network training feasible; however, it is biologically implausible and potentially interferes with topographic map formation, since changing input weights to a unit in one part of the cortical sheet will also change the weights of many other distant units in a non-local fashion. Some topographic models avoid this issue by forgoing the use of convolutional layers altogether, but in doing so forfeit the ability to model retinotopically-organized cortical areas. In contrast, our approach is to pre-optimize unit positions (see Methods) in a way that allows the learning of locally-smooth topographic maps even with convolutional layers (see Methods). In the brain, a similar pre-optimization may be achieved by chemical gradients [98] and experience-independent refinement of neural circuits during embryonic development[99, 100, 101, 102].

Finally, an exciting application of the TDANN is the simulation of experiments with spatial manipulations and readouts (Figure 8). Virtually every experiment that uses topographic structure as a dependent variable, including controlled rearing and task learning paradigms, could first prototype experiments with TDANNs. In addition, experiments that involve inactivation or stimulation of local populations of neurons (e.g. Rajalingham and DiCarlo [103], Shahbazi et al. [91]) could use the TDANN to predict the downstream behavioral impact of those manipulations prior to collecting data. The tools to perform stimulation or inactivation of neural populations have become commonplace in systems neuroscience in the past decade, but their engagement with the strongest models of neuronal function – task-optimized neural networks – has been limited due to the lack of image-computable models that not only explain the responses of individual neurons [104, 105, 106, 25] but that are also mapped to cortical tissue. As a unified model of functional organization, the TDANN is well-suited to bridge this gap.

## Methods

### Code and data availability

Code for model training and analyses is available at https://github.com/neuroailab/TDANN.

### Neural network architecture and training

#### Model training

We build off of the *torchvision* implementation of ResNet-18 [59] and train models with modifications to the VISSL framework [107]. All models were trained for 200 epochs of the ILSVRC-2012 (ImageNet Large-Scale Visual Recognition Challenge; Deng et al. [64]) training set. Unless otherwise indicated, models were each trained from five different random initial seeds. Network parameters were optimized with stochastic gradient descent with momentum (*γ* = 0.9), a batch size of 512, and a learning rate initialized to 0.6 then decaying according to a cosine learning schedule [108]. Models were trained either for supervised 1000-way object categorization or on the self-supervised contrastive objective “SimCLR” [63]. Following training, categorization accuracy for self-supervised models was assessed by freezing the parameters of the model and training a linear readout from the outputs of the final layer. The linear readout is trained for 28 epochs with a batch size of 1,024 and a learning rate initialized to 0.04 and decreasing by a factor of 10 every eight epochs.

#### Initialization of model unit positions

Prior to training, model units in each layer are assigned fixed positions in a two-dimensional cortical sheet that is specific to that layer. For efficiency, we do not embed the units of the very first convolutional layer. The size of the cortical sheet in each layer depends on a mapping between model layers and regions in the human ventral visual pathway, as well as a commitment to the extent of the visual field being modeled. For example, because we map model Layer 4 to human V1, the surface area of the cortical sheet in that layer is set to 13*cm*^2^: the mean value reported by Benson et al. [109] for the surface area of the section of human V1 that is sensitive to the central 7 degrees of visual angle. Another critical parameter in our framework is the size of a “cortical neighborhood”: during training, computation of the spatial loss is restricted to units within the same cortical neighborhood. We set the neighborhood width to match measurements made of the spatial extent of lateral connections in different cortical areas of the macaque (from Yoshioka et al. [110]), then scale up to achieve estimates that might match the human ventral visual pathway. Table 1 details the sizes of simulated cortical sheets and cortical neighborhoods in all layers.

**Table 1.** Parameters for layer positions. *the value of 1.6mm used in the V1-like layer is known to be inaccurate, but matching the proper value yields too few units in each cortical neighborhood to compute pairwise distances. See Supplementary Figure S5 for a solution to this problem.

| Layer | # Units | Size of Cortical sheet | Neighborhood Size | Region |
| --- | --- | --- | --- | --- |
| Layer 2 | 200704 | $5.7\text{mm}^2$ | $47\mu\text{m}$ | Retina |
| Layer 3 | 200704 | $5.7\text{mm}^2$ | $47\mu\text{m}$ | Retina |
| Layer 4 | 100352 | $13.5\text{cm}^2$ | $1.6\text{mm}^*$ | V1 |
| Layer 5 | 100352 | $13.5\text{cm}^2$ | $1.6\text{mm}^*$ | V1 |
| Layer 6 | 50176 | $12\text{cm}^2$ | $4\text{mm}$ | V2 |
| Layer 7 | 50176 | $5\text{cm}^2$ | $2.5\text{mm}$ | V4 |
| Layer 8 | 25088 | $49\text{cm}^2$ | $31\text{mm}$ | VTC |
| Layer 9 | 25088 | $49\text{cm}^2$ | $31\text{mm}$ | VTC |

### Positions are assigned in a two-stage process

#### Stage 1: Naive Retinotopic Initialization

Because each layer performs a convolution over the previous layer’s outputs, responses are organized into spatial grids. We preserve this intrinsic organization by assigning each model unit to a region of the simulated cortical sheet that corresponds to its spatial receptive field.

#### Stage 2: Pre-optimization of positions

Convolutional networks share filter weights between units at different locations; thus, local updates to a single unit entail updates to all units with the same filter weights. It is highly unlikely that an arbitrary configuration of unit positions will permit local smoothness under this global coordination constraint. Thus, we perform pre-optimization of unit positions to identify a set of unit positions for which learning smooth cortical maps is possible. Specifically, we spatially shuffle the units of a pre-trained DCNN on the cortical sheet such that nearby units have correlated responses to a set of sine grating images. The choice of sine gratings here is inspired by observations that edge-like propagating retinal waves drive experience-independent organization of the visual system in primates and other mammals [99, 100, 101, 102].

The spatial shuffling works as follows: 1) Select a cortical neighborhood at random. 2) Compute the pairwise response correlations of all units in the neighborhood. 3) Choose a random pair of units, and swap their locations in the cortical sheet. 4) If swapping positions decreases local correlations (measured as an increase in the Spatial Loss function described below), undo the swap. 5) Repeat steps 3-4 500 times. 6) Repeat steps 1-5 10,000 times.

#### Loss functions

We use two kinds of loss functions: spatial losses that encourage topographic structure, and task losses that encourage the learning of visual representations. We detail each in turn below:

##### Spatial loss

The spatial loss (SL) function encourages nearby pairs of units to have response profiles that are more correlated with one another than those of distant of units. Consider a neighborhood with *N* units. The vector of pairwise Pearson’s response correlations, 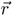, has length 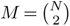, the number of unique pairs. Let the corresponding vector of pairwise Euclidean cortical distances be denoted 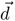.

We define two SL variants:

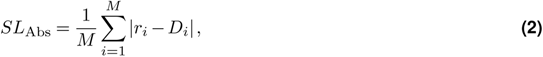

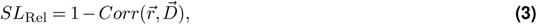

where *Corr* is the Pearson’s correlation function and 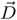 is the inverse distance:

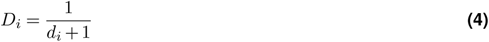

##### Task loss

The task loss is computed from the output of the final model layer. We use two task losses: the object categorization cross-entropy loss used in supervised object recognition (e.g. Krizhevsky et al. [111]) and the self-supervised SimCLR objective [63].

##### Combination of losses during training

On each batch, model weights are updated to minimize a weighted sum of the task loss and the spatial loss contributed by each layer:

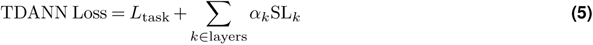

where *α* is the weight of the spatial loss.

##### Overview of Training

In summary, models are trained in 6 steps:

1. ResNet-18 is trained on the task loss only.
2. Positions in each layer are initialized to preserve coarse retinotopy (Stage 1).
3. Positions are further pre-optimized in an iterative process that preserves retinotopy while bringing together units with correlated responses to sine gratings images (Stage 2).
4. Positions are frozen and never again modified.
5. All network weights are randomly re-initialized.
6. The network is trained to minimize a weighted combination of the spatial and task loss components.

### Benchmarks comparing macaque V1 to model V1-like layers

#### Stimuli and Tuning Curves

##### Sine Grating Images

Tuning to low-level image properties such as orientation, spatial frequency, and chromaticity was assessed by constructing 224 *×* 224 pixel sine grating images that span 8 orientations evenly spaced between 0 and 180 degrees, 8 spatial frequencies between 0.5 and 12 cycles per degree, 5 spatial phases, and two chromaticities: black/white gratings and red/cyan gratings.

##### Tuning Curves

We evaluated tuning for orientations and spatial frequencies by constructing tuning curves for each unit. Color-responsiveness is assessed by comparing the mean response to all black and white gratings to the mean response to all red/cyan gratings. The distribution of model unit activations for a given layer was rescaled to match the minimum and maximum firing rates reported in [34]. We quantify the orientation tuning strength of model units using circular variance (CV), where values closer to 0 correspond to sharper tuning. As in Ringach et al. [34], CV is defined as:

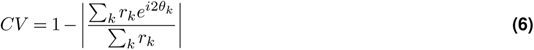

Where *θ_k_* is the *k*th orientation, in radians, and *r_k_* is the scaled response to that orientation. Orientation tuning curves are additionally fit with a von Mises function whose peak is taken as the preferred orientation.

#### Models

##### Hand-Crafted Self-Organizing Map

Our hand-crafted self-organizing map (SOM) implementation uses the *MiniSom* library [112], with parameters adapted from Swindale and Bauer [11]. We instantiate the SOM as a 128 x 128 grid of model units.

10,000 training samples were randomly constructed by selecting a random (x, y) location, orientation ([0, *π*], spatial frequency ([0, 1]), and chromaticity (black/white, colorful).

As in Swindale and Bauer [11], SOM weights were initialized retinotopically with randomly-selected initial preferred orientations.

The SOM is trained by presenting training examples for a total of 700,000 updates. After each example, the “winning” unit (i.e. the one with the highest response) is updated with a learning rate of ɛ = 0.02 to be more strongly aligned with the input stimulus, and its neighbors are updated in proportion to their proximity to the winner, as determined by a Gaussian neighborhood function parameterized by *σ* = 2.5.

Following training, each sine grating in the set of probe stimuli is presented to the SOM by projecting it into the six-dimensional space of SOM unit tuning and computing the response of each SOM unit to the stimulus. Once responses to each stimulus are obtained, tuning curves are constructed as usual.

##### DNN-SOM

The DNN-SOM is identical to the hand-crafted SOM, except that 1) the inputs are derived from the outputs of the first layer of an AlexNet model pretrained for ImageNet object categorization and 2) the learning rate is increased, which we found helps convergence. Following the approach of Zhang et al. [12], we take the responses of the first AlexNet layer to all 50,000 natural images in the ImageNet dataset, reduce their dimensionality with principal components analysis, and train the SOM on those examples.

#### Response Benchmarks

Model responses are compared to macaque V1 by considering preferred orientations and orientation tuning strength. Orientation tuning strength is computed as circular variance (CV) and compared between the population of model units and the empirical distribution provided by Ringach et al. [34] with the Kolmogorov-Smirnov distance. To filter out noisy units, we compute CV for model units with a mean response magnitude of at least 1.0. The distribution of preferred orientations is also compared to empirical data collected by De Valois et al. [35] by counting the number of units preferring each of four orientations: 0, 45, 90, and 135 degrees. In Figure S3b we compute a “Cardinality Index”: the fraction of preferred orientations that include, 0, 90, and 180 degrees.

#### Topographic Benchmarks

Orientation preference maps (OPMs) are compared to empirical measurements in two ways: counting pinwheels and quantifying map smoothness.

##### Pinwheel Detection

We interpolate the OPM onto a two-dimensional grid by computing the circular mean of the preferred orientation of units near a given location. If the population of model units near a grid location has high heterogeneity in preferred orientation, we disqualify that pixel for having an unreliable estimate of preferred orientation. Each grid location is assigned a “winding number” [17], computed by considering the preferred orientations of the eight pixels directly bordering the pixel under consideration. Moving clockwise around the bordering eight pixels, the change in preferred orientation from pixel to pixel is summed. A high winding number indicates a clockwise pinwheel, and a low winding number indicates a counterclockwise pinwheel, where the thresholds for “high” and “low” are selected to be consistent with manual annotation of clear pinwheels.

##### Pairwise Tuning Difference

We compute the smoothness of orientation preference maps by constructing a curve relating pairwise difference in preferred orientation to pairwise cortical distance. First, we restrict the population of model units to those with the highest 25% peak-to-peak tuning curve magnitudes. This filtering step removes units with weak responses or responses that would be indistinguishable from a “cocktail blank” background activity level, and we consider it equivalent to neuron selection in electrophysiological and optical imaging studies [34, 43]. As in similar approaches to quantifying OPM structure (e.g. Chang et al. [68]), pairs of units are binned according to their distance, and the average absolute different in preferred orientation is plotted for each distance bin. Because there can be hundreds of thousands of units in a given layer, we restrict this analysis to randomly-selected neighborhoods of a fixed width, then sample many neighborhoods from each map. Finally, we divide the pairwise distance by the chance value obtained by random resampling of unit pairs, such that a values < 1 indicate more similar tuning than would be expected by chance.

The OPM curves are compared to reconstructed macaque V1 data from Nauhaus et al. [43].

We adopt an identical approach for the construction of a neural spatial frequency preference map, where data are also provided for the same imaging window in Nauhaus et al. [43]. A similar strategy was used to recover data on cytochrome oxidase (CO) uptake from Livingstone and Hubel [38].

##### Smoothness

We define a smoothness score for a given map by comparing the tuning similarity for the nearest model unit pairs to the tuning similarity of the least similar pairs. Concretely, given a vector *x* of pairwise tuning similarity values, sorted in order of increasing cortical distance:

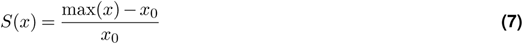

### Benchmarks comparing human VTC to model VTC-like layers

#### Stimuli

We evaluate the selectivity of neurons and model units to visual object categories using the “fLoc” functional localizer stimulus set [76]. fLoc contains five categories, each with two subcategories consisting of 144 images each. The categories are faces (adult and child faces), bodies (headless bodies and limbs), written characters (pseudowords and numbers), places (houses and corridors), and objects (string instruments and cars). Selectivity was assessed by computing the *t*-statistic over the set of functional localizer stimuli and defining a threshold above which units were considered selective.

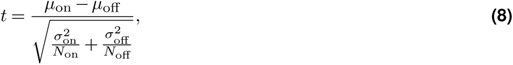

where *μ*_on_ and *μ*_off_ are the mean responses to the “on” categories (e.g., adult and child faces) and “off” categories (e.g., all non-face categories), respectively, *σ*^2^ are the associated variances of responses to exemplars from those categories, and *N* is the number of exemplars being averaged over.

#### Human Data

We compare models to human data from the Natural Scenes Dataset (NSD) [75], a high-resolution fMRI dataset of responses to 10,000 natural images in each of eight individuals (see Allen et al. for details). Models are compared to two aspects of this dataset: single-trial responses to the main set of natural images per participant (see “One-to-one mapping”) and selectivity in response to the “fLoc” stimuli. Single-trial responses were *z*-scored across images for each voxel and session and then averaged across three trial repeats. Selectivity was computed on the “fLoc” experiment as described in the previous section, generating *t*-maps for each of the five categories for each individual subject.

The VTC region of interest (ROI) was drawn based on anatomical landmarks to follow the convention in the literature [113] and is provided in the NSD data release as the “Ventral” ROI in the “streams” parcellation.

#### Models

##### Interactive Topographic Network (ITN)

We reconstruct maps from a variant of the ITN in Blauch et al. [20] that was trained and evaluated on the same images as the remaining models.

##### DNN-SOM

Two related approaches for building SOM models of higher visual cortex have recently been published [12, 13]. Because neither paper evaluates the resulting topographic maps with the fLoc stimuli, we reimplement the approach of Zhang et al. [12] as follows. We extract the responses of each unit in the final layer of a pretrained AlexNet to all 50,000 images in the ImageNet validation set. The responses are then reduced to the first four principal components. The SOM is initialized as a 200 x 200 grid of model units with a Gaussian neighborhood function set to *σ* = 6.2. The learning rate is set to 1.0 and the SOM is trained for 200,000 total iterations. The fLoc images are presented to the pretrained AlexNet model and projected into the space spanned by the four principal components computed previously. The response of each model unit to each fLoc image is computed by taking the dot product of the unit weight matrix with the projected fLoc images. The SOM is then treated identically to the VTC-like layer of TDANN.

#### Response Benchmarks

##### Representational similarity analysis

We compare functional properties of human VTC and models with representational similarity analysis (RSA) [72]. For any given model or human hemisphere, we compute a representational similarity matrix (RSM) as the pairwise Pearson’s correlation between patterns of selectivity for each of the five fLoc categories. The diagonal of the RSM is trivially 1.0 and is ignored in further analysis. The similarity of two RSMs is computed as Kendall’s *τ*.

#### Topographic Benchmarks

##### Pairwise Tuning Difference

We measure pairwise difference in VTC-like layer unit tuning as a function of cortical distance. We draw 25 randoms samples of 500 units each. Each sample is filtered to include only units with a mean response of at least 0.5 a.u.. For each fLoc category, the absolute pairwise difference in selectivity is computed for pairs of units separated by different cortical distances. Curves are normalized by the chance value obtained by randomly shuffling unit positions. Smoothness of maps is computed from these curves, same as in our analysis of V1. To compare a model to a human hemisphere, we compute the mean category-by-category difference in smoothness, e.g. comparing model face map smoothness to human face map smoothness, model body map smoothness to human body map smoothness, etc. Permutation tests randomly assigning category-by-category smoothness profiles to either “model” or “human” were used to assess the statistical significance of the mean difference in smoothness.

##### Patch Count and Size

Patches are automatically detected in maps of category selectivity by identifying contiguous regions of highly-selective units (or voxels, for human VTC). Patch identification has a small number of parameters that can be adjusted for maps of different sizes and with different dynamic ranges of selectivity values. The first step in identifying patches is to smooth and interpolate discrete selectivity maps. The selectivity map is then thresholded, and contiguous islands surviving the threshold are retained as candidate patches. Each candidate patch is further filtered for reasonable size: patches must be at least 100*mm*^2^ and no larger than 45*cm*^2^. Finally, the 2D geometry of the patch is constructed by fitting the concave hull of the points within the patch.

The following table identifies the relevant parameters for patch identification in human VTC and for each candidate model class.

**Table 2.**
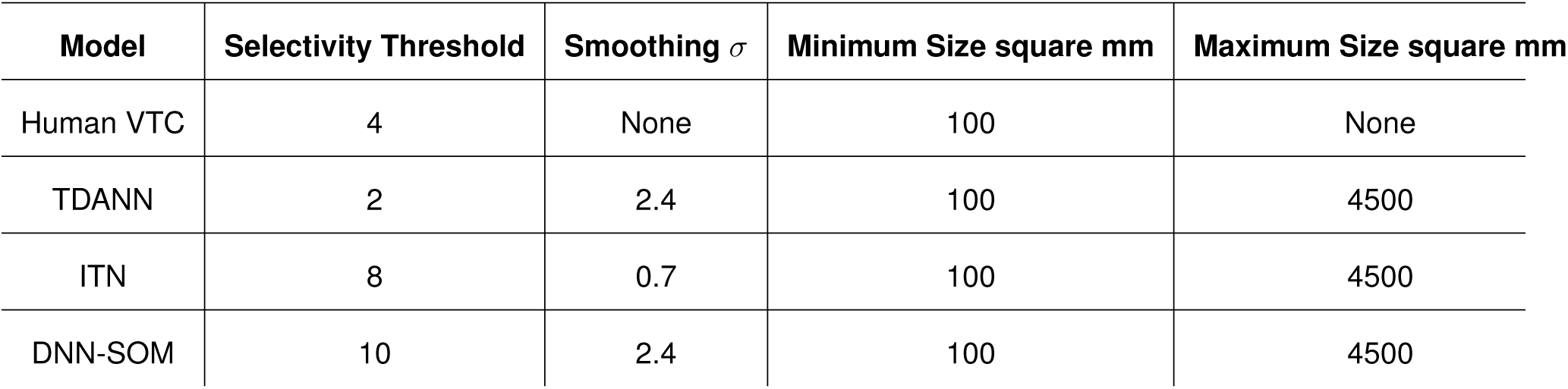
Patch detection parameters for human VTC and each model.

| Model | Selectivity Threshold | Smoothing $\sigma$ | Minimum Size square mm | Maximum Size square mm |
| --- | --- | --- | --- | --- |
| Human VTC | 4 | None | 100 | None |
| TDANN | 2 | 2.4 | 100 | 4500 |
| ITN | 8 | 0.7 | 100 | 4500 |
| DNN-SOM | 10 | 2.4 | 100 | 4500 |

##### Selectivity Overlap

We determine if units (or voxels, for human VTC) that are selective for a pair of categories overlap with one another as follows. First, we bin the cortical sheet into discrete square neighborhoods of width 10mm. In each neighborhood, the fraction of units selective for Category X and Category Y are recorded. We consider two populations as overlapping if there is a strong correlation between the proportions recorded across neighborhoods, i.e., if the frequency of Category 1 selectivity is predictive of Category Y selectivity and vice-a-versa. The X-Y Overlap score is computed as

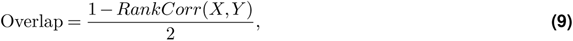

where RankCorr is the Spearman’s rank correlation coefficient and 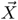 is the proportion of units selective for Category X in each cortical neighborhood. The category selectivity threshold was set at *t* > 4.

#### Linear regression

Neural predictivity is computed against a given dataset as the mean variance explained across neurons and splits of the data. In practice we follow the parameters and design decisions made by the BrainScore team [30]; they are repeated here for completeness. We use partial least squares (PLS) regression to predict the activity of a given neuron as a linear weighted sum of model units in a given layer. Model activations are preprocessed by first projecting unit responses to ImageNet images onto the first 1000 principal components, i.e. each component is a linear mixture of model units. This projection is used when fitting on the stimuli that were shown to the animal. When fitting IT, we use data from Majaj, Hong, et al., 2015 [32], which consists of multi-electrode array data in responses to quasi-naturalistic scenes with a variety of objects on a variety of backgrounds. Variance explained is corrected by dividing raw predictivity by the internal noise ceiling, a measure of the consistency of each recorded neuron.

### One-to-one mapping of visual cortical responses

A direct, one-to-one mapping between units and voxels is computed by assigning each unit in a layer of the network to a single voxel based on responses to a given dataset. In practice, we correlate individual model unit activations to the natural images from the Natural Scenes Dataset [75] with responses to these same images on the single voxel level for a given subject. Unit-to-voxel assignments are determined using a polynomial-time optimal assignment algorithm [114] which maximizes the overall average correlation between unit and voxel pairs, on a given training set. The 515 shared images that all eight subjects viewed three times were held out as a test set and all reported one-to-one correlations are calculated on this test set, using the unit-to-voxel assignments determined from training. Each unit-to-voxel correlation is normalized by the individual voxel noise ceiling of that assigned voxel (see Allen et al. for information on the calculation of the intra-individual voxel noise ceilings in NSD). One-to-one correlations were calculated on an individual subject basis for each of the self-supervised and supervised models trained at each level of the spatial weight *α*. The inter-individual, or subject-to-subject, noise ceiling, was calculated in the same manner, this time assigning voxels from one subject to voxels from another subject based on how correlated responses to the shared 515 images were for each potential voxel pair. For the subject-to-subject assignment, we used an 80/20 train/test split and averaged results for each subject combination across 5 splits. A similar analysis will appear in a forthcoming publication by Finzi et al.

### Wiring Length

We measure the functional wiring length between two adjacent layers, the “source” layer and the “target” layer by first identifying the units with the highest responses in each layer, then computing the length of inter-layer fibers that would be required to connect them. First, for a given natural image input, we identify the top *p*% most responsive units in each of two adjacent layers. We set *p* to 5% in the V1-like layers and 1% in the VTC-like layers. We note that for computational tractability, we restrict our analysis to small neighborhoods in the V1-like layers and average results across many random neighborhood selections.

Next, inter-layer fibers are added one by one, until all activated units in the earlier “source” layer are sufficiently close to the location at which a fiber originates. In practice, we find the optimal fiber origination sites using the *k*-means clustering algorithm, and continue adding fibers until the total “inertia” of the *k*-means clustering falls below a specified threshold, *k*_thresh_. Inertia is computed as the sum of the squared distances between each activated unit and its nearest fiber, and *k*_thresh_ is set such that the mean distance from each unit to its nearest fiber is not greater than *d*_thresh_. *d*_thresh_ is set to 10.0mm in the VTC-like layer pairs, and is reduced to 0.9mm in the V1-like layer pairs to reflect the smaller cortical neighborhood. Having established the number of inter-layer fibers required and their origination sites in the “source” layer, we identify optimal termination sites for those fibers in the “target” layer as follows. The set of target layer termination sites is identified as the centroids from *k*-means clustering, with *k* set to the number of fibers. Finally, fibers are assigned between origination sites and termination sites with the linear sum assignment algorithm, and the total wiring length is computed as the sum of the lengths of each individual inter-layer fiber.

A critical decision when measuring wiring length in this way is how to situate units from two layers in a common physical space. By design, each TDANN layer occupies a unique two-dimensional sheet, leaving the spatial relationships between units in different cortical sheets undefined. Here, we assume that the “source” cortical sheet and “target” cortical sheet lie in the same 2D plane, joined at one edge. Concretely, we can position the “target” sheet to the left, right, above, or below the “source” layer. Without reason to choose one of these strategies, we compute the optimal wiring length for each of the four options and report the average across all shift directions.

### Dimensionality

In our analyses of dimensionality, we consider the responses of the full population of model units in each layer to a set of 10,112 natural images from the NSD [75]. Following [83], we perform spatial max-pooling on the convolutional feature maps, then compute the eigenspectrum of these responses. We summarize the dimensionality of the responses by their effective dimensionality (ED; Del Giudice [84]):

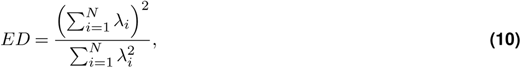

where *λ_i_*is the *i*th eigenvalue, and *N* is the number of eigenvectors.

### Microstimulation of model units on the simulated cortical sheet

We simulate the microstimulation of local populations of model units to 1) gain insight into the functional properties of local populations, and 2) measure effective connectivity between groups of units in adjacent layers. In all analyses, stimulation is performed by fixing the activity of units to values determined by a 2D Gaussian function. Units near the center of the Gaussian have their activity set to the maximal value, and activity falls off with distance from the center. We consider the top 5% of units, ranked by activity level, as being responsive in either the “Source” layer, where activity is set according to the 2D Gaussian, or in the following “Target” layer, where unit activity is determined by the network architecture and learned weights.

#### Functional Alignment

In VTC-like layers, we measure functional alignment between layers by comparing the category selectivity of activated units in the Source layer (Layer 8) with the selectivity of responsive units in the Target layer (Layer 9). For each stimulation site, we compute the mean selectivity (*t*-statistic) of the top 5% most activated units for each of the following categories: faces, bodies, characters, cars, and places. This five-element “selectivity profile” can then be compared to the profile of the top 5% most strongly responding units in the Target layer by computing *χ*^2^ distance between selectivity profiles. Similarity is then taken as the negative log distance and compared to a shuffle-control in which a random subset of units is compared instead of the top 5% most active units.

### Simulation of a Visual Cortical Prosthesis

In Figure 8, we demonstrate a proof of concept for using topographic DCNNs to prototype visual cortical prosthetic devices. This proof of concept consists of two distinct stages: 1) generating device-achievable stimulation patterns with a Stimulation Simulator, and 2) generating the estimated percept (Percept Synthesizer) that would result by stimulating cortical areas with those patterns. To generate stimulation patterns, we feed a target image into TDANN and record the precise activation magnitude of each model unit in each layer. If an infinitely high-precision stimulation device with absolute coverage of the cortical sheet in all cortical areas were available, we would stimulate cortex with this set of precise activation patterns. However, real stimulation devices are limited in many ways, including limits to their spatial precision and the set of cortical areas they can access. Thus, we use TDANN to produce *device-achievable* stimulation patterns, i.e., those that are consistent with the limitations of cortical stimulation devices. Here we take a simple approach by considering degradation of high-precision patterns into device-achievable patterns by Gaussian blurring. In each layer, we first interpolate the precise activity patterns onto a high-resolution grid (2500 *×* 2500 px), then blur the resulting pattern with a 2D Gaussian kernel whose *σ* parameter is set according to the desired blur level. Because different layers have different cortical sheet sizes (e.g. 70mm on an edge in the VTC-like layer and 37mm on an edge in the V1-like layer), the width of the Gaussian in *pixels* is variable, even though the width of the Gaussian in *mm* is constant. Finally, we perform a nearest-neighbor lookup such that each model unit adopts the activity level of the pixel closest to its location. This set of activity patterns is the final “device-achievable” pattern. The Stimulation Simulator also allows any specific subset of layers to be included; e.g. the first two layers only, or all eight layers. We consider this restriction comparable to the limited access a neural stimulation device might be restricted to.

Given a set of device-achievable activity patterns, we seek to determine the estimated percept that would be evoked if that pattern were written into cortex, i.e., the visual input that is most consistent with those patterns. To this end, we follow the example of Granley et al. [90] and use gradient-ascent image optimization methods to synthesize an image such that the activity pattern produced by presenting that image is as close as possible to the device-achievable target pattern. We use the *lucent* Python package to iteratively optimize an image to minimize the total mean squared error, summed across layers, between the target activity patterns and the current evoked patterns at that iteration. We optimize the image for 3000 steps at a learning rate of 0.05; further optimization has little effect on reducing the mean squared error. The optimized result is the predicted percept for a given input image and theoretical cortical stimulation device.

## Author Contributions

E.M. and D.F. performed analyses. E.M., K.G.-S., and D.L.K.Y. wrote the paper. H.L., J.J.D., and D.L.K.Y. originally conceived the approach.

## Acknowledgements

This work was supported by a National Science Foundation Graduate Research Fellowship awarded to E.M., a National Institutes of Health grant (RO1 EY 022318) awarded to K.G.-S., a Simons Foundation grant (543061) awarded to D.L.K.Y., a National Science Foundation CAREER grant (1844724) awarded to D.L.K.Y., and an Office of Naval Research grant (S5122) awarded to D.L.K.Y. We also thank the NVIDIA corporation and the Google TPU Research Cloud group for hardware grants. We are grateful to Ben Sorscher for helpful discussions.

## Supplementary Information

### V1-like maps produced with alternative feature sets

Figure 2 demonstrates that co-training for spatial and task losses is sufficient to generate V1-like topography. However, we have not ruled out the possibility that generating orientation-selective units and arranging them on the cortical sheet via other strategies could produce V1-like maps. To address this concern, we derive orientation prefrence maps (OPMs) from three different strategies for learning and spatially organizing model units. We first compare the standard TDANN, in which unit positions are fixed prior to training and model weights are optimized to minimize both task and spatial losses, to a Task Only DCNN whose weights are optimized only for the task loss. To generate an OPM from the Task Only model, we freeze network weights then iteratively shuffle model units on the cortical sheet such that the Spatial Loss is minimized post-hoc. Accordingly, we refer to this model as a “Post-hoc” arrangement of DCNN features. We find that OPM smoothness is nearly identical when co-learning features with the spatial loss (i.e., TDANN) than when first learning features and then post-hoc arranging units in the cortical sheet (Supplementary Figure S4). A third alternative is to bypass the learning of features altogether and use a hard-coded Gabor filterbank (GFB) to generate model units, as has been suggested as a model of V1 neuron tuning (e.g. Jones and Palmer [5], Dapello et al. [3]). Following the same approach as in the Task Only model for deriving OPMs, we find that the hard-coded GFB features fail to produce a smooth OPM. How can we reconcile the apparent inadequacy of the Gabor filterbank in generating V1-like topography with its strong orientation selectivity? One possibility is that a Gabor filterbank lacks the required complexity to form responses to natural images that drive brain-like topography, but that simpler stimuli may improve the accuracy of its topographic predictions. To test whether the nature of the images presented to the model matters, we evaluated the same three feature sets (TDANN, Post-hoc, Task Only, and GFB) on a set of simple sine grating images (Figure S4a, bottom). Interestingly, we find that the TDANN, Post-hoc Task Only, and GFB feature sets all produce smooth OPMs when their units are organized with respect to correlations of sine grating responses. We conclude that the TDANN is the only model that, by co-learning features and topography, is able to produce brain-like OPMs from realistically complex natural inputs. Task Only and Hand-Crafted feature spaces are capable of producing V1-like OPMs only when presented with simple inputs, whereas the core advantage of the TDANN is its ability to learn a feature space that produces brain-like functional organization in the presence of realistically complex natural images.

### Natural image inputs are required for the emergence of brain-like functional organization

Work in developmental neuroscience and psychology has called into consideration the influence of visual experience on the development of structure and function in visual cortex. We leveraged the ability of self-supervised TDANNs to predict functional organization after learning from unlabeled visual data streams to determine which inputs might drive the emergence of brain-like topographic maps. We evaluated networks trained on four distinct image datasets, including the natural image datasets ImageNet and Ecoset [9], and two artificial datasets: sine gratings and white noise images. We find that for both natural image datasets, there is brain-like functional organization of V1-like and VTC-like layers. In the V1-like layer, 14% of units in the Ecoset-trained network and 20% of units in the ImageNet-trained network were strongly orientation selective (circular variance < 0.6), and we observe smooth OPMs with pinwheels in models trained from both datasets. Further, we found similar numbers of VTC-like layer units with selectivity *t* > 5 in both models (12.7% for Ecoset and 14.2% for ImageNet), and we detect patches selective for all five categories in both models (Figure S9).

While the suitability of naturalistic stimuli for generating brain-like functional organization may not be surprising, we wanted to test if simpler artificial datasets could succeed in matching neural data for two reasons. First, it has been demonstrated that patterned endogenous activity prior to eye opening can establish visual cortical circuitry [4]. Second, if artificial synthetic stimuli were suitable for constructing brain models, we could avoid needing to collect large natural image datasets. We trained TDANN on two artificial stimulus sets: a set of sine grating stimuli that may loosely mirror endogenous activity patterns, and Gaussian white noise images. The grating-trained model exhibited a very high fraction of strongly orientation selective units in the V1-like layer (73%). However, the grating-trained model had no category selectivity (1.2% of units selective at *t* > 5, averaged across categories), and no detectable patches. Thus, simple oriented stimuli may be sufficient to drive V1-like map formation, but natural stimuli are necessary to develop the remainder of the ventral visual pathway. We next evaluated a model trained on white noise, which allows us to isolate the effects of the model architecture and loss functions in the absence of structure in the input data. We find that training on white noise prevents the learning of strongly orientation-selective units in the V1-like layer (0% of units with circular variance < 0.6) or strongly category-selective units in the VTC-like layer (4% of units with *t* > 5). Surprisingly, however, the noise-trained TDANN does learn some weak functional organization. In the V1-like layer, a weak orientation preference map is formed, and in the VTC-like layer, two character-selective patches and one face-selective patch is observed. These results suggest that the spatial loss is able to produce some topographic structure even in the absence of patterned inputs, although the strength of the selectivity is extremely weak. Taken together, these analyses of the impact of training data on functional organization support the necessity and sufficiency of natural images for the emergence of robust V1-like and VTC-like topographic maps.

### Probing the tuning of unit populations outside of category-selective patches

If the VTC-like layer smoothly encodes a space of objects, we might expect that images synthesized to drive high responses in nearby regions of the cortical sheet would be perceptually similar. Indeed, we find that optimal image characteristics smoothly vary across the cortical surface. Higher spatial frequency and rectilinear features dominate the upper right sides of the map, while curvilinear and lower spatial frequency features best drive the top and bottom edges. We find that these optimal images also align with category-selectivity, e.g., the input images that best drive units in face-selective patches tend to contain eyes and fall in the more curvilinear regions of feature space. Images synthesized to maximize regions that fall between patches (sites 5, 10, 11, 15, 16, and 20) lack clearly discernible object categories, but nonetheless follow the smooth gradients of image features across the cortical surface. Thus, it appears that the VTC-like layer learns a smooth mapping of object space in two dimensions, and that patches emerge as regions of that space that align with the category localization stimuli that we use to probe the model.

### Supplemental Methods

#### Dimensionality Summarize by Power Law Exponent

Following Kong et al. [7], Stringer et al. [11], we summarize the eigenvalues by fitting a line to the log-log plot of eigenvalues against their principal component index, and report the absolute value of the best fit line as the power law exponent. To prevent fitting to nonlinear regions in the earliest and latest parts of the distribution, the line is fit from the 2nd to the 50th principal component.

#### Linear regression

Neural predictivity is computed against a given dataset as the mean variance explained across neurons and splits of the data. In practice we follow the parameters and design decisions made by the BrainScore team [10]; they are repeated here for completeness. We use partial least squares (PLS) regression to predict the activity of a given neuron as a linear weighted sum of model units in a given layer. Model activations are preprocessed by first projecting unit responses to ImageNet images onto the first 1000 principal components, i.e. each component is a linear mixture of model units. This projection is used when fitting on the stimuli that were shown to the animal. When fitting V1, we use data from Cadena et al. [2], which consists of single-neuron recordings to a set of natural images. When fitting V4 and IT, we use data from Majaj, Hong, et al., 2015 [8], which consists of multi-electrode array data in responses to quasi-naturalistic scenes with a variety of objects on a variety of backgrounds. Variance explained is corrected by dividing raw predictivity by the internal noise ceiling, a measure of the consistency of each recorded neuron.

#### Unit Clustering

The degree to which of responses to natural images are clustered is computed by considering the locations of the 5% of units that respond most strongly to a given input image. We compute the distribution of pairwise distances between these highly-active units, then count the number of pairs that are within 10.0mm of each other: if the count is high, then the active units are concentrated into a small number of clusters. Finally, clusterness is defined as the ratio between the number of nearby pairs in the true response pattern to the number of nearby pairs when locations are randomly shuffled. We compute results for 10 random position shuffles, 64 randomly-selected images used as input, and five random initial seeds for each model.

#### Gabor Filter Bank (GFB)

In Figure S4, we generate responses from a Gabor filter bank by following the VOneNet implementation in Dapello et al. [3]. For computational tractability and to produce a similar quantity of units as in the TDANN V1-like layer, we reduce the number of simple and complex channels from 256 to 64 each, and increase the stride of the convolution from 4 to 8 pixels. The resulting filter bank is then treated identically to the TDANN V1-like layer when extracting responses and constructing tuning curves.

The orientation preference map (OPM) for the GFB model is produced by assigning GFB outputs to random initial positions, then minimizing the Spatial Loss by iteratively swapping the locations of randomly-selected pairs of units as described above.

#### Stimulus Optimization

We use image synthesis methods, implemented in the *lucent* Python package (https://github.com/greentfrapp/lucent), to generate images which reproduce patterns of stimulation. Specifically, we synthesize an input image that minimizes the mean squared error between a desired pattern of activity and the actual pattern obtained by presenting the synthesized image to the network. The desired pattern of activity is set according to a two-dimensional Gaussian centered over some region of the cortical sheet. In these experiments we set the *σ* parameter of the Gaussian to 3.5mm. For efficiency, we also remove units far from the center of the Gaussian from the computation of the mean squared error: units below 10% of the height of the Gaussian are ignored. All synthesized images begin as 128 x 128 pixels of white noise and are optimized for 1,024 steps. We retain the default settings for image transforms, which include optimization in the Fourier basis, color channel decorrelation, jittering, rotation, and padding.

**Figure S1.**
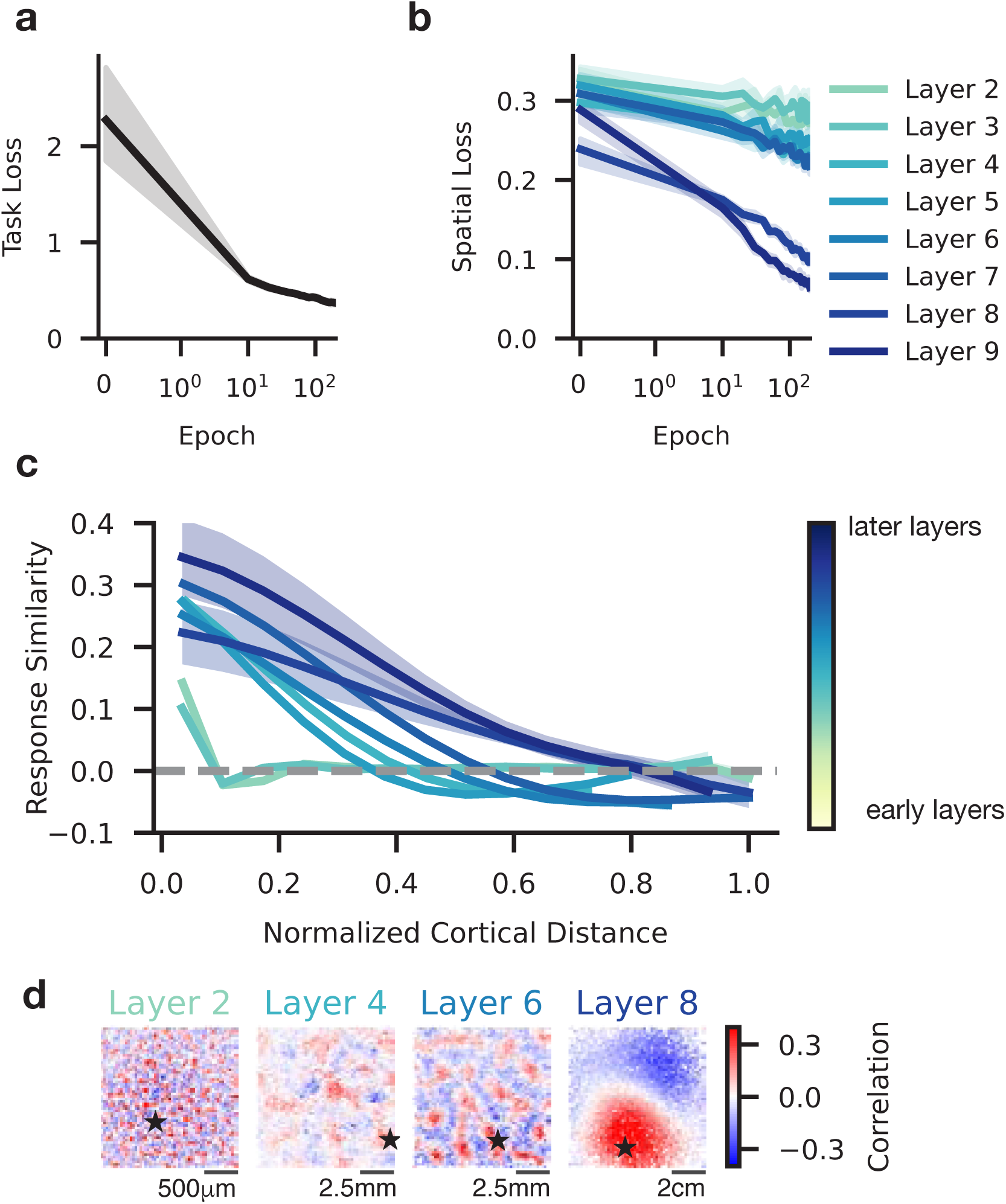
Minimization of loss components during training. **(a)** Task loss throughout training. **(b)** Spatial loss in each of the eight convolutional layers during training. Shaded area: 95% confidence interval (CI) across random initializations. **(c)** Response correlation decreases as a function of the cortical distance between model unit pairs in each model layer. Shaded region: 95% CI from repeated sampling of different cortical neighborhoods in each layer. **(d)** Portions of the cortical sheet from each of four convolutional layers; units colored according to their correlation with an arbitrarily selected seed unit, marked by the black star.

#### Retinal Waves

In Figure S18 we organize DCNN units in the cortical sheet according to their response correlations to a series of simulated retinal waves.

#### Creating Retinal Wave

Our simulation of retinal wave activity is heavily inspired by the description in Kim et al. [6]. We simulate the retina as a two-dimensional circle of radius 320px. The retina has three spatially-overlapping cell layers: one for ON-RGCs (retinal ganglion cells), one for OFF-RGCs, and one for amacrine cells. Each cell can be in one of four states: inhibited, recruitable (but not currently active), refractory (recently active but not recruitable yet), or active (currently “on”). Cells are connected to each other according to the following rules: 1) ON-RGCs are connected to one another in an excitatory fashion within a radius of *r*_on_, 2) ON-RGCs are connected to amacrine cells in an excitatory fashion within a radius of *r*_on_, and amacrine cells inhibit OFF-RGCs within a radius of *r*_amacrine_.

**Figure S2.**
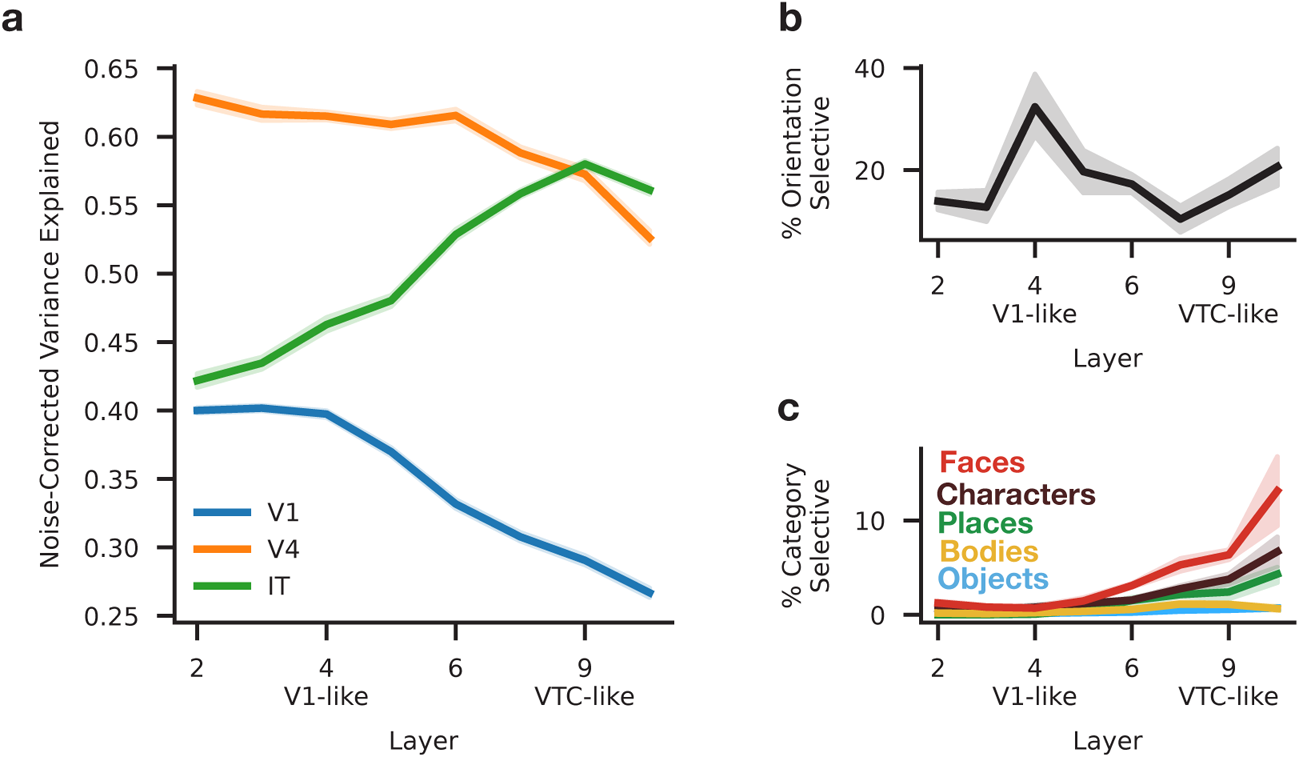
Selection of V1-like and VTC-like layers. **(a)** Variance explained by linear regression of TDANN layer outputs to measurements in macaque V1, V4, and IT. V1 predictivity peaks in the first three layers, whereas IT predictivity peaks in the last two layers. **(b)** Fraction of units strongly orientation selective in each layer. **(c)** Fraction of units that are strongly selective for each category (*t*-value > 10) in each layer.

**Figure S3.**
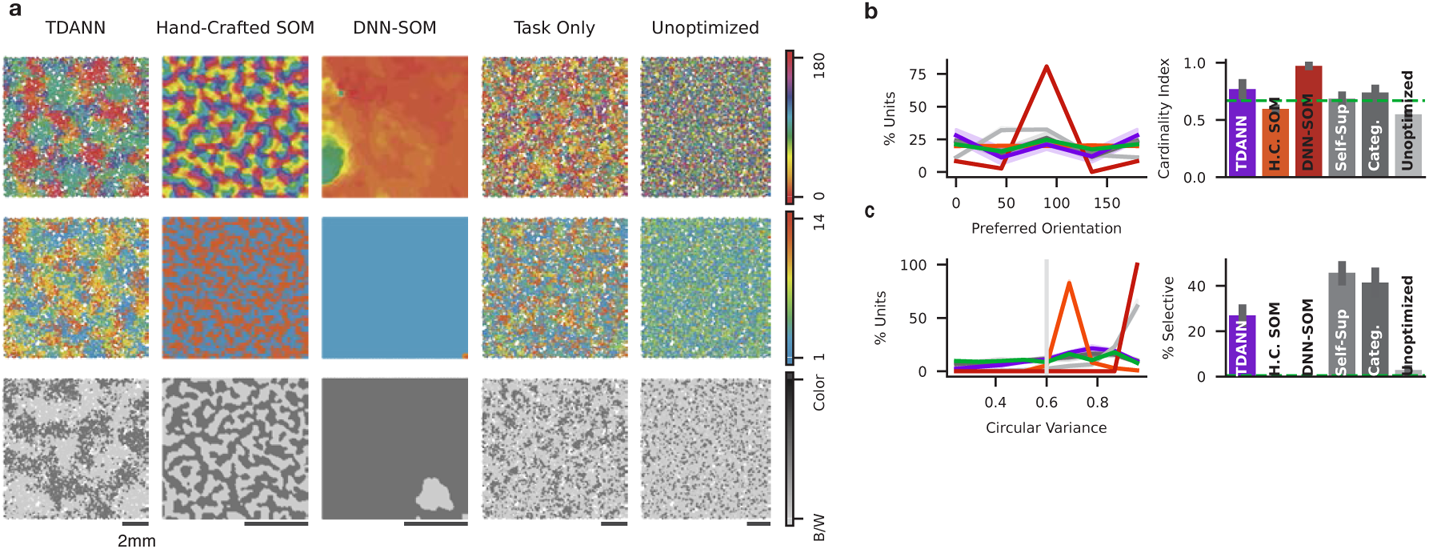
Topographic and representational benchmarks in the V1-like model layer. **(a)** Orientation, spatial frequency, and chromatic preference maps for all candidate model types. **(b)** Left: Distribution of preferred orientations for each model type. Right: Cardinality index, computed as the fraction of units selective for cardinal orientations to units selective for the obliques. Dashed green light indicates value in macaque V1. **(c)** Left: Distribution of circular variance for each model and for macaque V1. Vertical line indicates cutoff for strong selectivity. Right: percentage of units strongly selective for orientations in each model type.

A wave is initiated by setting some subset of the ON-RGCs to the “active” state. The activated subset is determined by picking a random location along the edge of the retina and activating cells along a thin strip at that location. The wave is then propagated for up to *t* timesteps (propagation is halted if the wave runs off screen and all cells are off). At each timestep, activity is propagated as follows. First, all cells that have been active longer than a specified “active duration” are set to the refractory state. Second, cell activity levels are updated by multiplying the connectivity matrices with the previous activity states. Third, we activate all ON-RGCs who are in the recruitable state and whose activity exceeds an activity threshold of *t*_ON-active_. Fourth, we inhibit all OFF-RGCs whose activity falls below a threshold of *t*_OFF-active_. OFF-RGCs whose activity passes that threshold are activated if they are currently in the recruitable state. Finally, amacrine cells whose activity exceeds *t*_amacrine-active_ are set to active. The remainder of the amacrine cells are made recruitable instead. Images of the simulated activity at each stage are produced by creating binary masks of the locations of active ON-RGCs. Half of the waves are randomly assigned to map the binary images to a black and white colormap, and the remainder are assigned to a red and green colormap.

**Figure S4.**
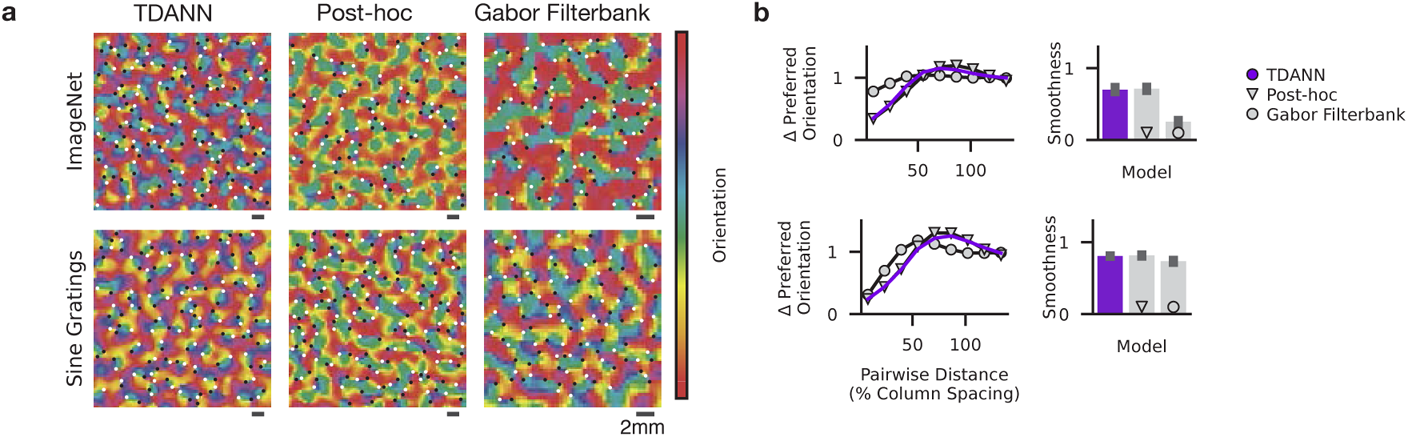
Smoothed OPMs from alternative feature spaces. **(a)** Top row: smoothed OPMs from the TDANN, a Task Only model with post-hoc unit organization, and a Gabor Filterbank with post-hoc unit organization, where units are brought closer together if they have similar responses to ImageNet images. Bottom row: same as top, but with unit proximity optimized with respect to sine grating image responses. **(b)** Pairwise orientation tuning difference over distance, and corresponding smoothness scores, for the maps in (a).

**Figure S5.**
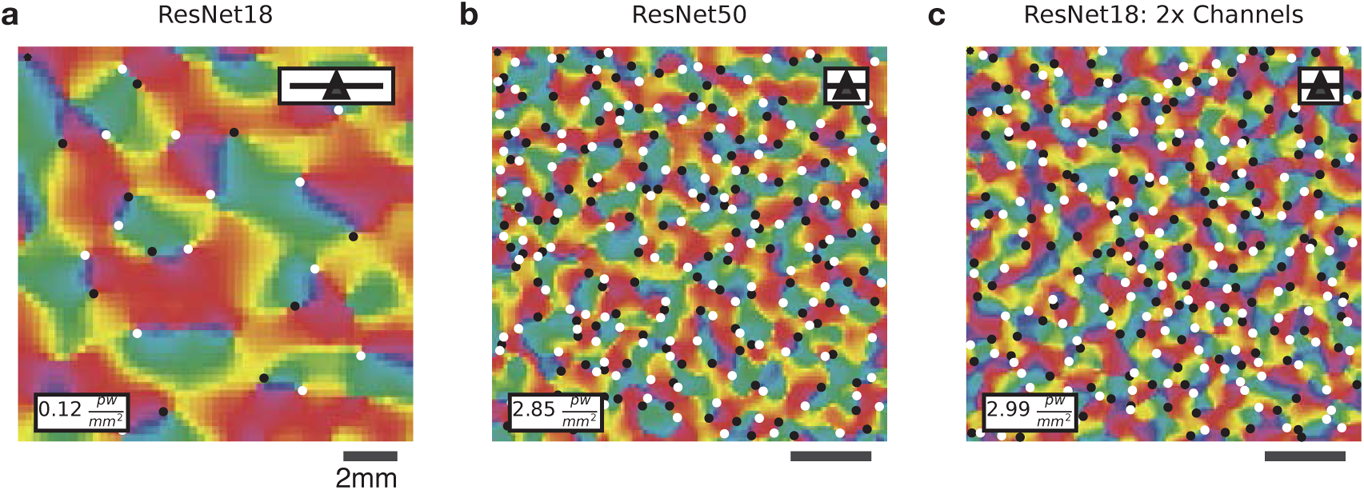
Orientation preference maps (OPMs) and pinwheel density in alternative models. For demonstration, all models in this figure had unit positions organized post-hoc to achieve a strong OPM, i.e., they are not proper TDANNs. **(a)** OPM in a small region of the standard ResNet-18 TDANN. Pinwheels are shown by black and white dots. **(b)** OPM in the V1-like layer of a categorization-trained ResNet-50, in which the increased number of channels allows a reduction of cortical neighborhood size and, accordingly, a dramatic increase in pinwheel density. **(c)** OPM in the V1-like layer of a categorization-trained ResNet-18 with twice the number of channels in each layer, in which the increased number of channels allows a reduction of cortical neighborhood size and, accordingly, a dramatic increase in pinwheel density.

In this work, we produce retinal waves with two sets of parameters. The following parameters are common to both sets of retinal waves: *r*_on_ = 15, *r*_amacrine_ = 1.5, *t* = 20, *t*_ON-active_ = 7, *t*_amacrine-active_ = 0.1, *t*_OFF-active_ = 0.1. In one of the two sets of waves, the active duration is set to 100ms, and in the other, the active duration is set to 200ms. In practice, the waves produced with the longer active duration are twice as thick.

#### Measuring Responses to Waves

Each wave consists of a number of images, one per timestep of the simulation. Because the simulated retina is circular, the corners of each image never contain simulated activity. To make better use of each image, we take a central square crop of each image of size M x M pixels then resize the image back to 224 x 224 pixels. *M* is selected such that all regions of the crop contain activity: for an image of size 224px, 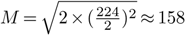. As with all other images presented to the DCNN models, the images are then preprocessed and normalized.

**Figure S6.**
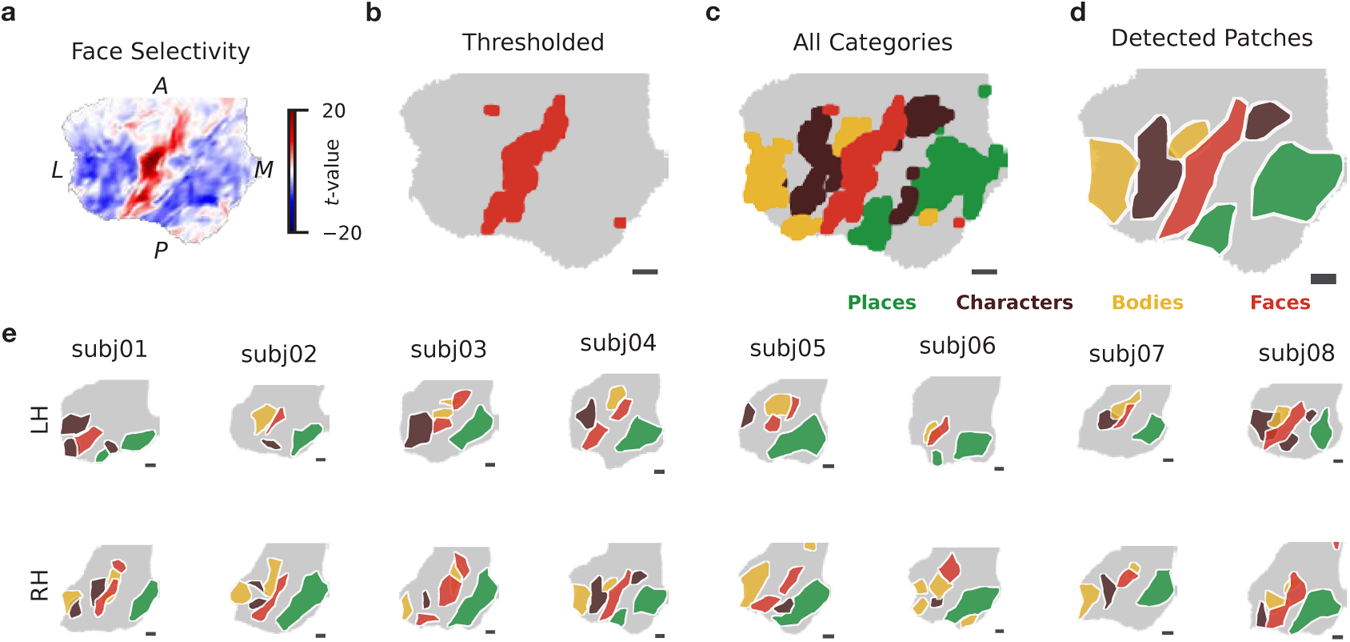
Data in each human subject from the NSD fLoc experiment and patch detection protocol. All scale bars: 2mm. **(a)** Map of face selectivity in the right-hemisphere VTC region of interest (ROI) for one example subject. A: anterior, M: medial, L: lateral, P: posterior. **(b)** Thresholded face selectivity map for the same subject as (a). **(c)** Category selectivity map for all five fLoc categories. **(d)** Patches detected from the category selective clusters in (c). **(e)** Detected patches in each hemisphere (LH = left hemisphere, RH = right hemisphere) for each subject.

**Figure S7.**
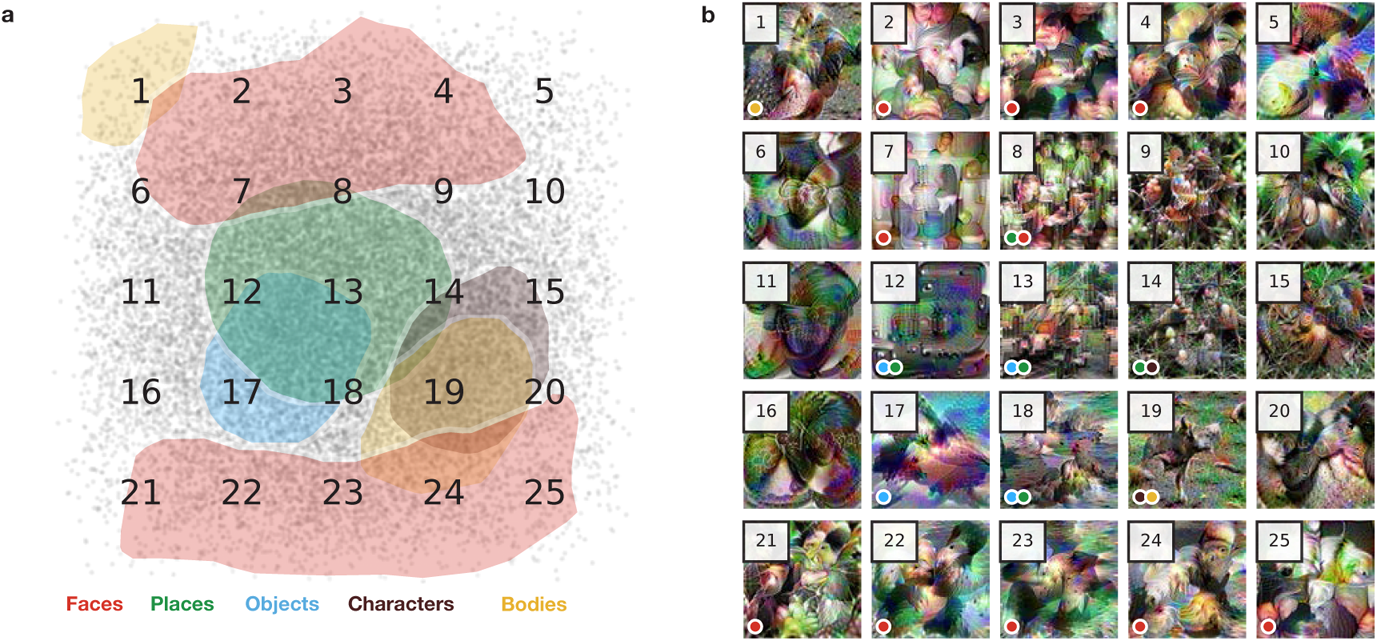
Optimal stimuli throughout the VTC-like layer. **(a)** VTC-like layer of an example TDANN model. Overlaid numbers correspond to sub-panels in (b). **(b)** Images synthesized to maximally activate a local population of units centered at the indicated location in (a). Small dot in bottom left of each image indicates the patch membership of that location, e.g., a red dot indicates that the image optimally drives units that happen to be in a face-selective patch.

For a wave with *t* timesteps, each model unit produces *t* responses. We integrate responses to each wave by computing the mean response across all waves. Anecdotally, similar results are achieved by computing the maximum response instead of the mean. Unit-to-unit correlations are then computed by considering the vector of integrated responses for each wave. We use the unit-to-unit correlations to perform swap-based organization of units on the cortical surface such that correlated units are moved to be nearby one another.

**Figure S8.**
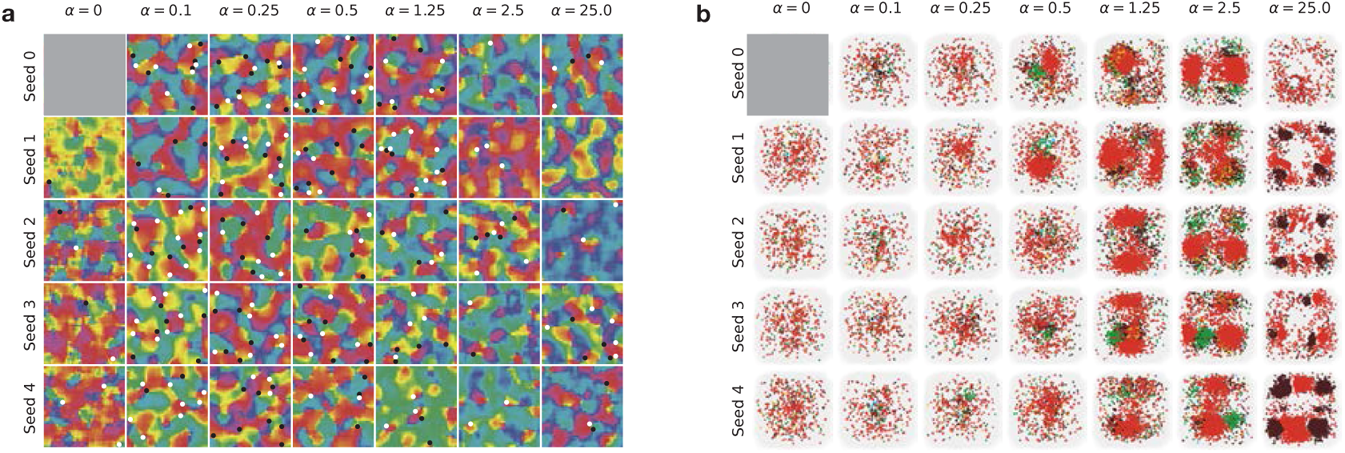
Topographic maps for models trained with the Relative SL and the Supervised Categorization objective. **(a)** Orientation preference maps (OPMs) in the V1-like layer of models at each level of *α* trained from five different random seeds with the categorization objective. A region of each cortical sheet is shown, with black and white dots indicating locations of detected clockwise and counter-clockwise pinwheels, respectively. Gray square covers the Task Only seed 0, which was used during position initialization. **(b)** Category selectivity maps for the VTC-like layer of each model in (A). Plotting conventions as in Figure 3.

**Figure S9.**
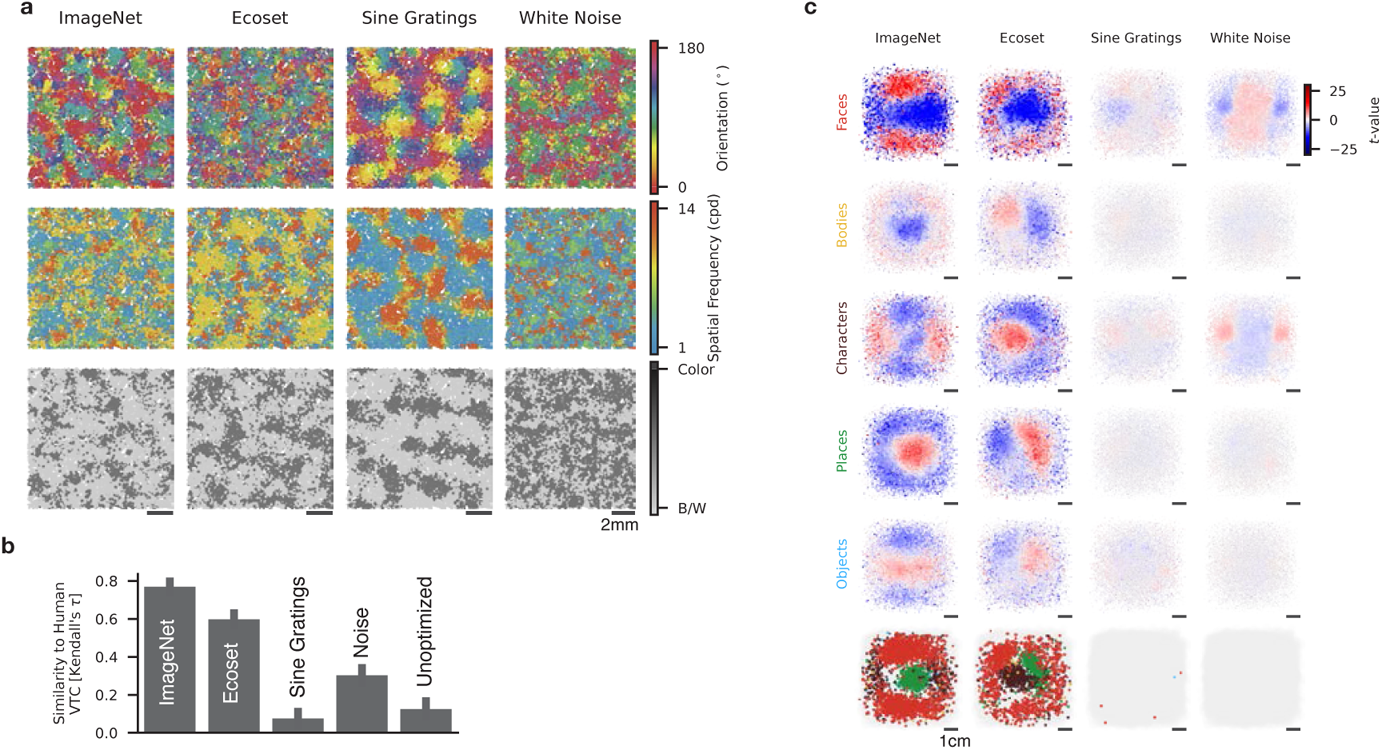
Topographic maps from models trained with different training datasets. **(a)** Orientation, spatial frequency, and chromatic preference maps in the V1-like layer of models trained with ImageNet images, the *Ecoset* training set, a set of hand-selected sine gratings (increased *α* = 10), and Gaussian white noise images. **(b)** Representational similarity between human VTC and models trained with each dataset. Error bar: 95% CI across human hemispheres. **(c)** Category selectivity maps for the VTC-like layer of each model in (a). Plotting conventions as in Figure 3.

**Figure S10.**
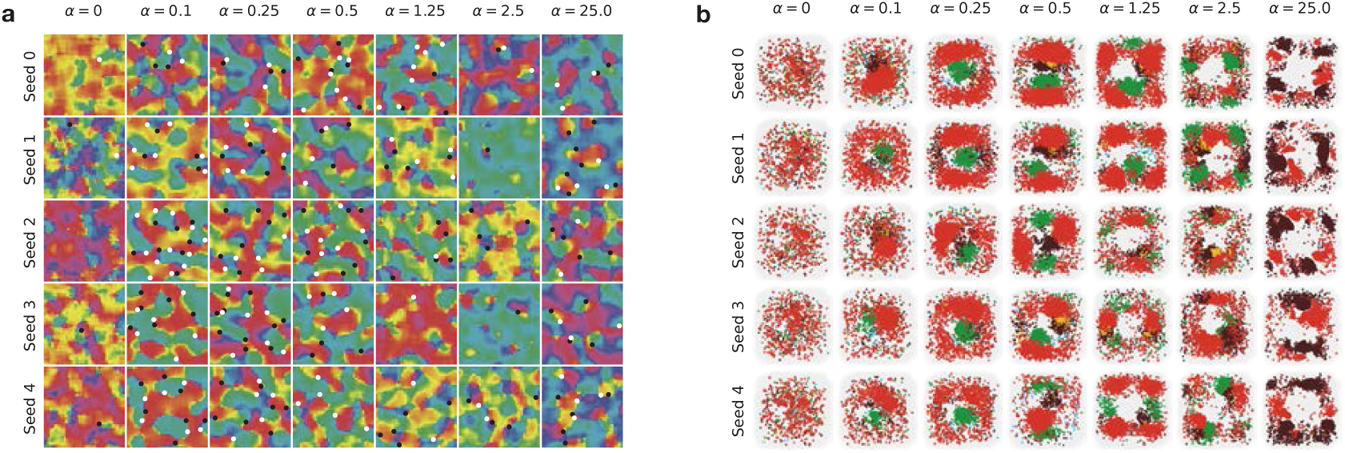
Topographic maps for models trained with the Relative SL and self-supervision. **(a)** Orientation preference maps (OPMs) in the V1-like layer of models at each level of *α* trained from five different random seeds with the Relative Spatial Loss (SL). A region of each cortical sheet is shown, with black and white dots indicating locations of detected clockwise and counter-clockwise pinwheels, respectively. **(b)** Category selectivity maps for the VTC-like layer of each model in (a). Plotting conventions as in Figure 3.

**Figure S11.**
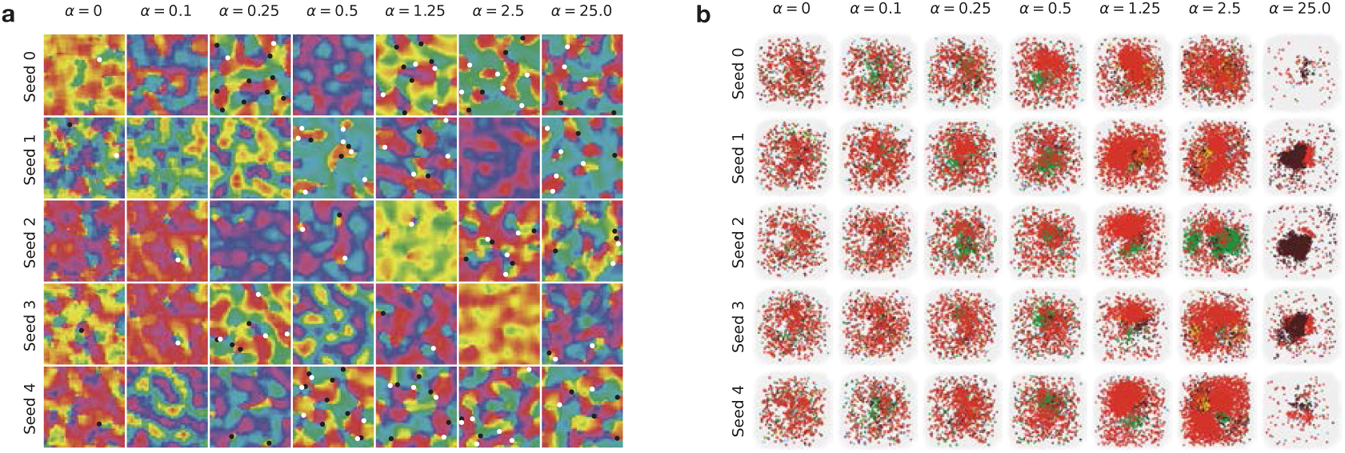
Topographic maps for models trained with the Absolute SL. **(a)** Orientation preference maps (OPMs) in the V1-like layer of models at each level of *α* trained from five different random seeds with the Absolute Spatial Loss (SL). A mm region of each cortical sheet is shown, with black and white dots indicating locations of detected clockwise and counter-clockwise pinwheels, respectively. **(b)** Category selectivity maps for the VTC-like layer of each model in (a). Plotting conventions as in Figure 3.

**Figure S12.**
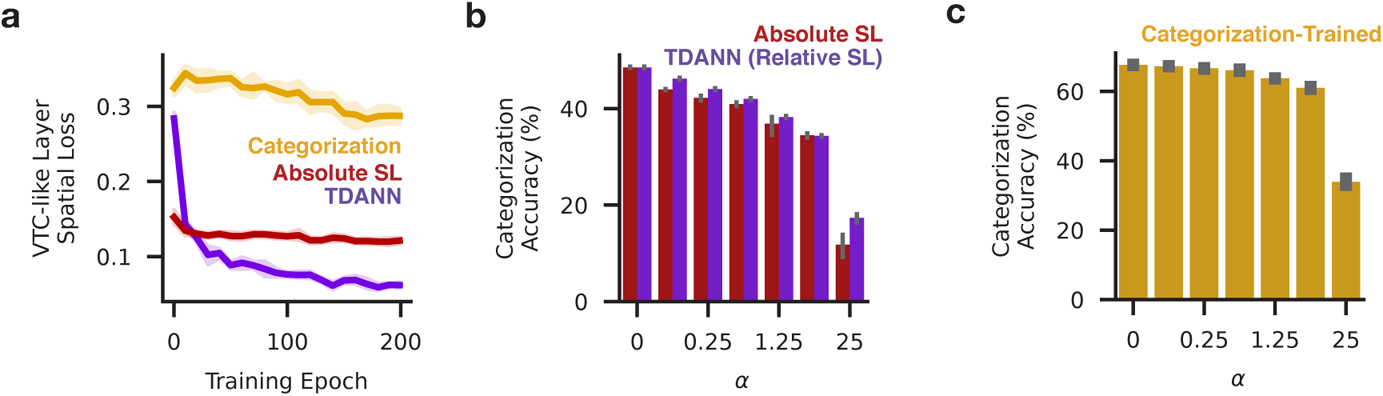
Additional comparison of models trained with different task and spatial objectives. **(a)** Spatial loss in the VTC-like layer of TDANN models (purple), categorization-trained models (gold), and models trained with the Absolute SL (red) throughout training. **(b)** Categorization accuracy (top-1 ImageNet validation set performance) for models trained at each level of *α* with either the Relative (purple) or Absolute (red) SL. **(c)** Categorization accuracy for models trained at each level of *α* directly on the supervised categorization objective.

**Figure S13.**
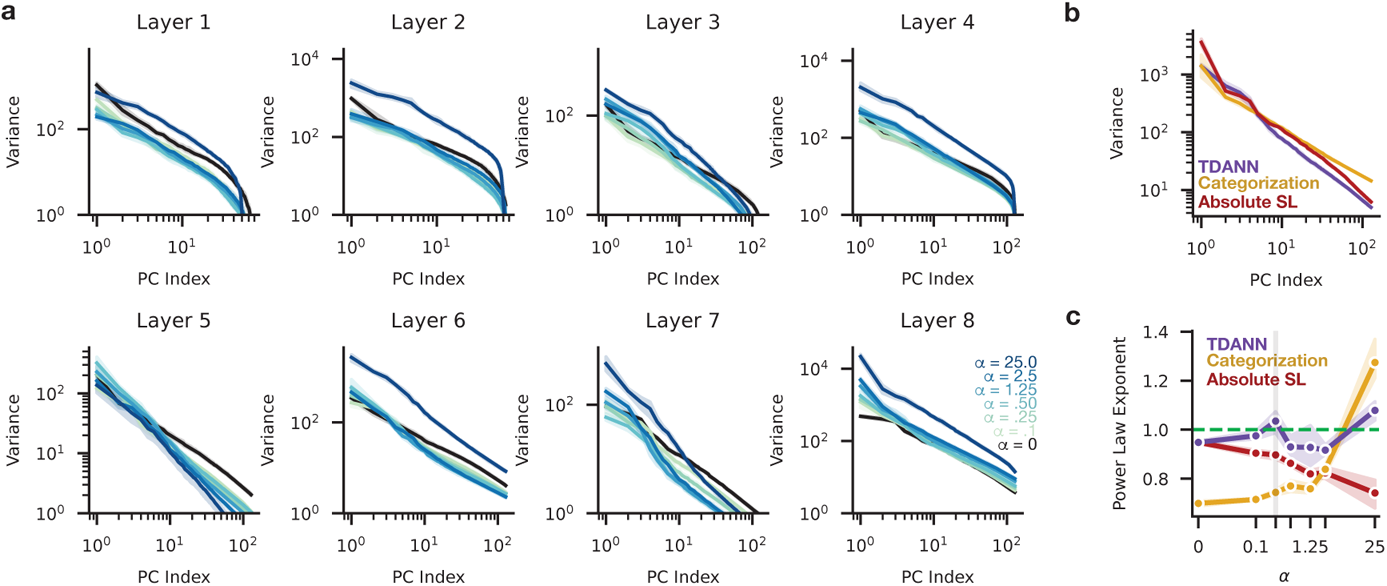
Dimensionality of model unit populations as a function of training objective and *α*. **(a)** Variance explained by each principal component (PC) for each layer of TDANNs trained at different levels of the spatial weight magnitude *α*. Components computed from responses to 10,000 images from the NSD [1]. **(b)** Variance explained by each principal component in the VTC-like layer of models trained with *α* = 0.25 and different objectives. **(c)** Power law coefficient fit to eigenspectra from the VTC-like layer of models trained with *α* = 0.25 and different objectives.

**Figure S14.**
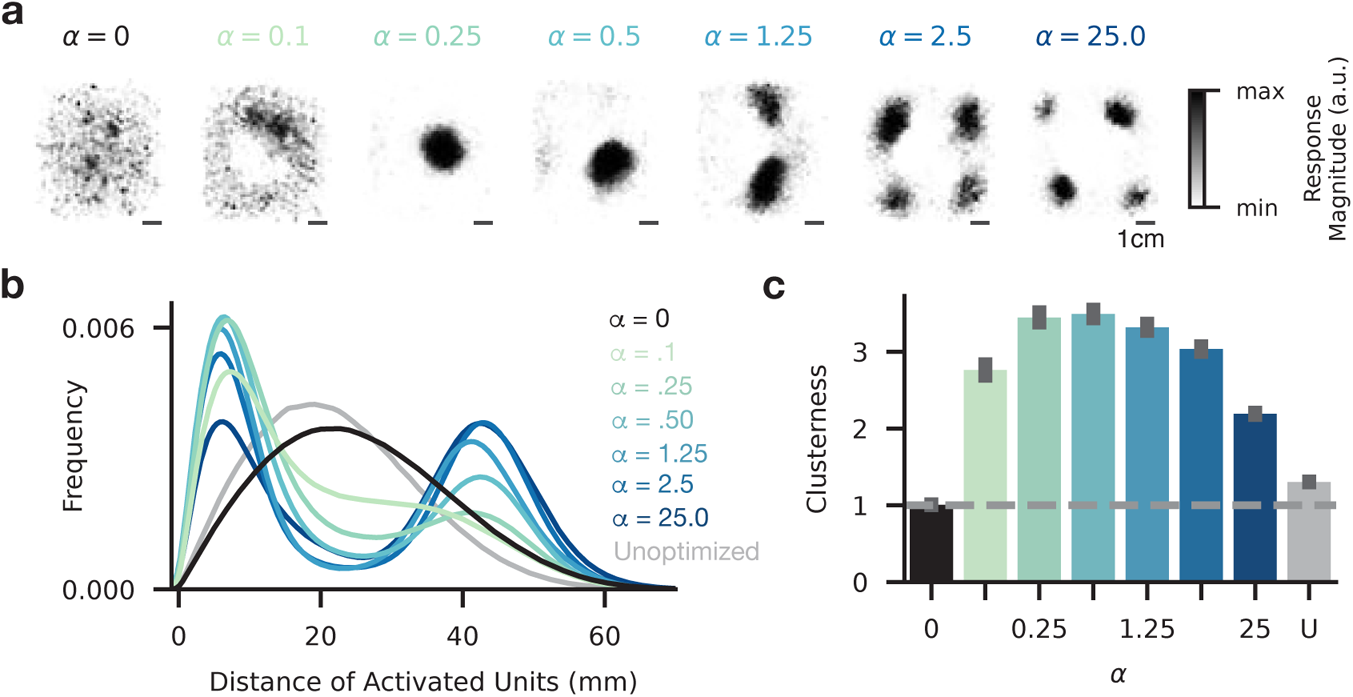
Clustering of responses to natural images as a function of *α*. **(a)** Strength of activation in the TDANN VTC-like layer to an arbitrarily-selected natural image, for models trained at different levels of the spatial weight (*α*). **(b)** Probability density function of pairwise distances between pairs of activated units for each model type, computed over repeated presentations of different natural images. Curve color indicates the level of the spatial weight (*α*) that model was trained with. **(c)** Clusterness, measured as the increase in unit density above the chance of value (dashed line: 1.0). Error bars: 95% CI over different random initial model seeds and images used to generate responses. ANOVA: *F* (7, 32) = 70.5, *p* < 10^−16^, post-hoc Tukey’s tests: significantly lower clusterness for *α* = 0 and Unoptimized models compared to models with *α >* 0, all post-hoc *ps* < .001.

**Figure S15.**
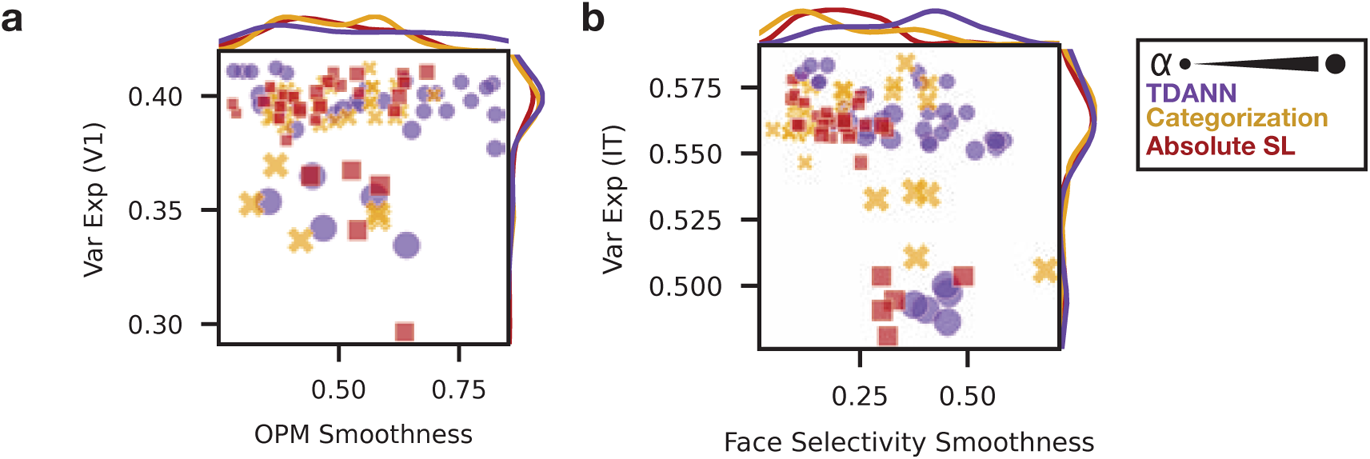
Prediction of neural firing rates with linear regression compared against topographic map smoothness. **(a)** Models trained at different levels of *α* (represented by dot size) and with different objectives compared in their capacity to predict macaque V1 firing rates (Var Exp) and the smoothness of their orientation preference maps. **b)** As in (b), but for prediction of firing rates in macaque inferotemporal cortex (IT) and smoothness of face selectivity maps. No difference in variance explained when *α* < 25, all pairwise *ps* from Mann-Whitney tests *p >* 0.42.

**Figure S16.**
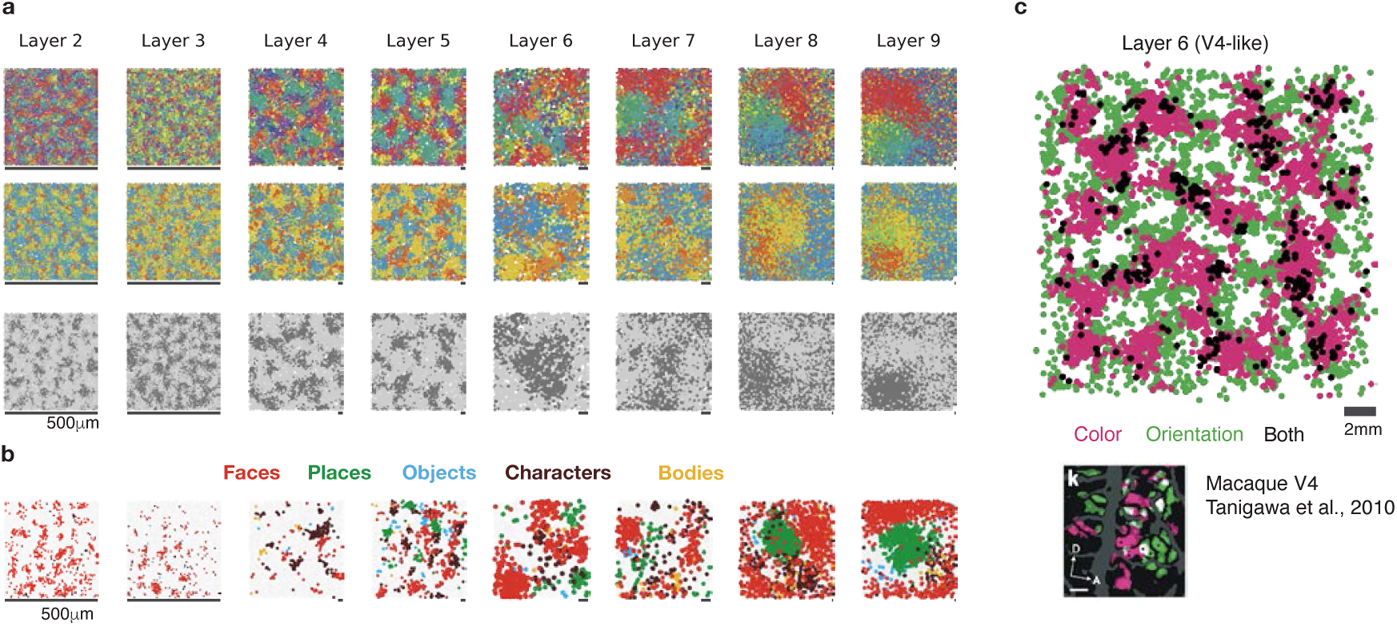
Topographic maps in each layer of a representative TDANN model. **(a)** Orientation, spatial frequency, and chromatic preference maps in each layer. Plotting conventions as in Figure 2. **(b)** Category selectivity map in each layer. **(c)** Orientation and color selectivity in the V4-like model layer. Units in magenta are selective for color and not orientation, units in green are selective for orientation and not color, and units in black are selective for both orientation and color. Similar data in macaque V4 is shown in the inset at bottom right (from [12]).

**Figure S17.**
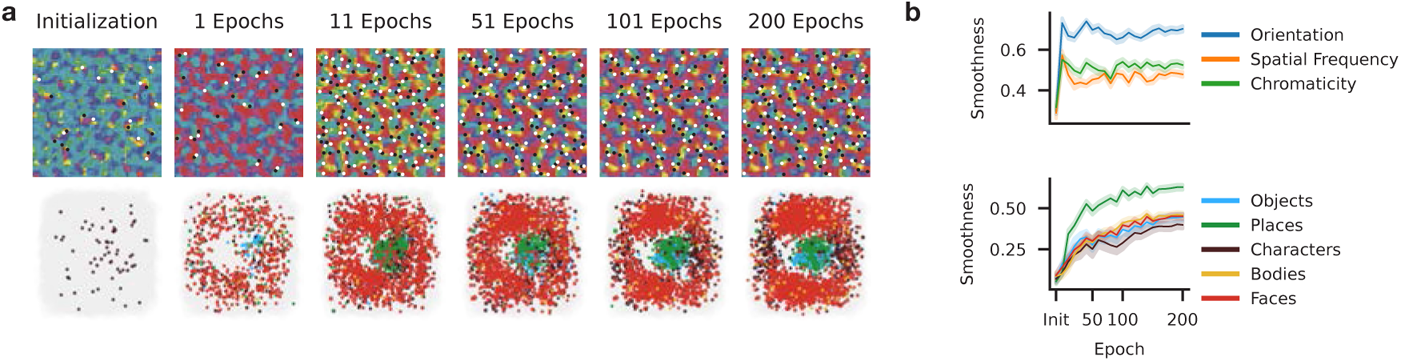
Topographic maps in a representative TDANN throughout training. **(a)** OPMs in the V1-like layer at initialization (left), and after 1, 11, 51, 101, and 200 epochs of training. **(b)** Category selectivity maps in the VTC-like model layer at each timepoint. **(c)** Smoothness as a function of training step for orientation, spatial frequency, and color preference maps. Smoothness peaks early then plateaus. **(d)** Selectivity of category selectivity maps for each fLoc category. Smoothness increases throughout training.

**Figure S18.**
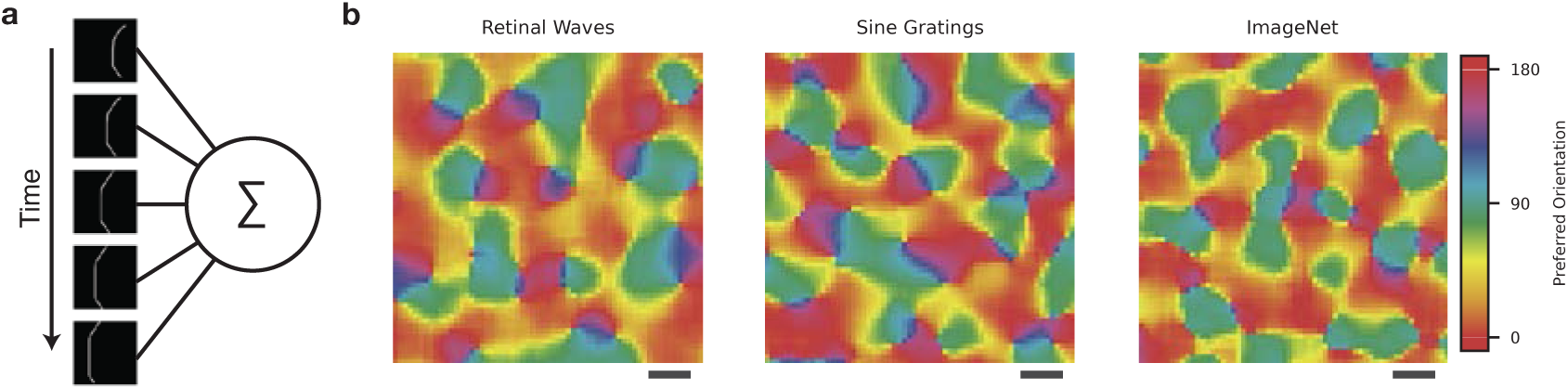
Simulated retinal waves can drive unit-to-unit correlations comparable to static sine gratings. **(a)** Five example frames from a simulated retinal wave movie. The responses to each frame are integrated to compute the mean response to each wave. **(b)** OPMs created by post-hoc organization of units in the V1-like layer of a Task Only SimCLR model, when the unit-to-unit correlations are computed by presenting retinal wave movies (left), a dataset of sine gratings (middle), or natural images (right). Scale bar: 2mm.

